# PYCR1-dependent proline synthesis in cancer-associated fibroblasts is required for the deposition of pro-tumorigenic extracellular matrix

**DOI:** 10.1101/2020.05.30.125237

**Authors:** Emily J Kay, Karla Paterson, Carla Riero Domingo, David Sumpton, Henry Daebritz, Saverio Tardito, Claudia Boldrini, Juan R Hernandez-Fernaud, Dimitris Athineos, Sandeep Dhayade, Ekaterina Stepanova, Enio Gjerga, Lisa J Neilson, Sergio Lilla, Ann Hedley, Grigorios Koulouras, Grace McGregor, Craig Jamieson, Radia Marie Johnson, Morag Park, Kristina Kirschner, Crispin Miller, Jurre J Kamphorst, Fabricio Loayza-Puch, Julio Saez-Rodriguez, Massimiliano Mazzone, Karen Blyth, Michele Zagnoni, Sara Zanivan

## Abstract

TElevated production of collagen-rich extracellular matrix (ECM) is a hallmark of cancer associated fibroblasts (CAFs) and a central driver of cancer aggressiveness. How to target ECM production to oppose cancer is yet unclear, since targeting CAFs has been shown to restrain but also promote cancer progression. Metabolic rewiring is a hallmark of CAFs. Here we find that proline, which is a highly abundant amino acid in collagen proteins, is newly synthesised from glutamine to make tumour collagen in breast cancer xenografts, and that its production is elevated in breast cancer CAFs. PYCR1 is the rate-limiting enzyme for proline synthesis and is highly expressed in the tumour stroma of breast cancer patients and in CAFs. Reducing PYCR1 levels in CAFs is sufficient to reduce tumour collagen production, tumour growth and metastatic spread in vivo and cancer cell proliferation in vitro. PYCR1 and COL1A1 are overexpressed in patients with invasive ductal carcinoma with poor prognosis. Both collagen and glutamine-derived proline synthesis in CAFs are enhanced by increased pyruvate dehydrogenase-derived acetyl-CoA levels, via gene expression regulation through the epigenetic regulator histone acetyl transferase EP300. Altogether, our work unveils unprecedented roles of CAF metabolism to support pro-tumorigenic collagen production. PYCR1 is a recognised cancer cell vulnerability and potential target for therapy, hence, our work provides evidence that targeting PYCR1 in tumours may have the additional benefit of halting the production of pro-tumorigenic ECM.

## Introduction

Cancer-associated fibroblasts (CAFs) are mesenchymal cells abundant in the stroma of solid tumours and active players in tumour initiation, progression, metastasis, and response to standard anti-cancer therapies (i.e. chemo-and radiotherapies) as well as targeted therapies (e.g. anti-angiogenic drugs) and immune therapies ^1–5^. Because of the extensive body of evidence that CAFs support cancer, targeting CAFs has emerged as a novel opportunity to control cancer. However, CAFs have also been shown to restrain tumour growth ^6, 7^. Therefore, a better understanding of CAF biology is a prerequisite in order to target them effectively to oppose cancer.

Distinct functional subpopulations of CAFs coexist in the stroma of solid tumours ^8–11^, and recent works suggest that they are dynamically interconvertible ^12, 13^. Among those, myofibroblast-like CAF (myCAF) resemble myofibroblasts ^14^ in that they produce abundant collagen-rich extracellular matrix (ECM) and express high levels of alpha smooth muscle actin (ACTA2, also known as αSMA) ^8–10^. The presence of myCAF in the tumour stroma has been described in human and murine solid cancers ^8, 10, 11, 15^. In patients, αSMA-expressing CAF abundance is an indicator of poor prognosis in several cancer types ^16–18^, including breast cancer where it has been associated with high histological grade, lymph node metastasis, high microvessel density, distant relapse and immunotherapy resistance ^9, 19–21^. Moreover, CAFs isolated from patients and kept in culture retain myCAF features ^9, 22, 23^ and when co-transplanted with cancer cells accelerate tumour growth and progression ^23–26^. Conversely, depletion of proliferating αSMA-expressing CAFs in murine breast cancer and PDAC models ^6, 27^ or the inhibition of their expansion through targeting the hedgehog pathway ^28^ in a PDAC model ^7^ accelerated cancer and promoted metastasis. Hence, controlling cancer progression may not require CAFs to be killed, but rather to target specific molecules and pathways that control their pro-tumorigenic functions.

The possibility of targeting ECM production for therapeutic purposes has received a lot of attention in the cancer field^29–33^, because the composition and mechanical properties of the ECM are established active drivers of tumour pathology ^34–37^. In particular, collagen, which is the most abundant component of the tumour ECM, has been associated with tumour-promoting functions. Genetic depletion of collagen VI (*Col6a1*) reduced the rate of tumour initiation and growth, while overexpression of collagen I (*Col1a1*) in breast cancer models has shown to enhance tumour formation and progression, and increase incidence of metastasis ^38–40^. Collagen also drives the formation of desmoplastic stroma, thus contributing to impeding effective delivery of therapeutics, as well as intratumoural recruitment of immune cells, by hampering the growth of a functional tumour vasculature ^29–33^. It is important to note that high-density non-fibrillar collagen I can protect against breast cancer development^41^, that the presence of fibrosis following neoadjuvant therapy in pancreatic cancer patients correlates with better prognosis ^42–44^ and that reducing tumour collagen production by deleting *Col1a1* in αSMA expressing cells or upon treatment with anti–lysyl oxidase like-2 antibody accelerated progression of pancreatic cancer ^45, 46^, suggesting that a collagen-rich ECM may have also tumour-protecting functions. This ambiguity underlines the need to understand how ECM production can be targeted effectively to oppose cancer. Several strategies have been developed to directly or indirectly target TGFβ signalling ^29–33^, a major trigger of fibroblast activation, which have shown synergistic effects with other anti-cancer therapies in pre-clinical models. However, TGFβ is a pleiotropic factor with both pro-and anti-tumorigenic functions ^47^, possibly explaining why only a handful of drugs have made it to the clinic and even fewer have provided a benefit to the patients ^3, 48, 49^. Hence, we need a better understanding of the molecular mechanisms that underpin the production of pro-tumorigenic ECM to develop complementary and different approaches that target ECM production in cancer.

Metabolic reprogramming is another feature of CAF activation that has been shown to have tumour promoting functions, primarily through providing tumour cells with nutrients, including lactate, pyruvate, glutamine, alanine, proline and ketone bodies ^50–58^. However, how to effectively target CAF metabolism in tumours is still an open question, since the metabolic vulnerabilities of CAFs and cancer cells can be very different, and crosstalk between the cell types creates an intertwined metabolic network. A deeper understanding of CAF metabolic reprogramming is needed to identify potential metabolic vulnerabilities to target CAF activation and tumour progression.

Using a previously characterised model of tumour-promoting mammary CAFs ^22, 25, 59^ and patient-derived CAFs isolated from aggressive breast tumours and their matched normal fibroblasts isolated from tumour-adjacent tissue (NFs), we show that the production of pro-tumorigenic collagens requires increased proline synthesis from glutamine. Moreover, we show how the rate limiting factor for proline synthesis, pyrroline-5-carboxylate reductase 1 (PYCR1), acetyl-CoA levels and the epigenetic regulator histone acetyl transferase EP300 are major regulators of this process.

## Results

### Myofibroblast-like CAFs use proline newly synthesised from glutamine for collagen production *in vitro* and *in vivo*

The production of abundant collagen-rich ECM is a trait that CAFs acquire during the transition from normal to activated fibroblasts and a major regulator of tumour progression ^1–4^. Amino acid frequency analysis of the matrisome ^60^ and of the total cell proteome pinpointed an exceptionally high content of proline and glycine residues in collagen proteins (**Figure 1a** and **Supplementary Data S1**). This is in line with the common knowledge that collagen proteins contain repeating glycine-proline-hydroxyproline sequences to be able to fold and assemble into stable fibres ^61^, and it raises the question of how CAFs can metabolically support this increased biosynthetic demand.

**Figure 1.**
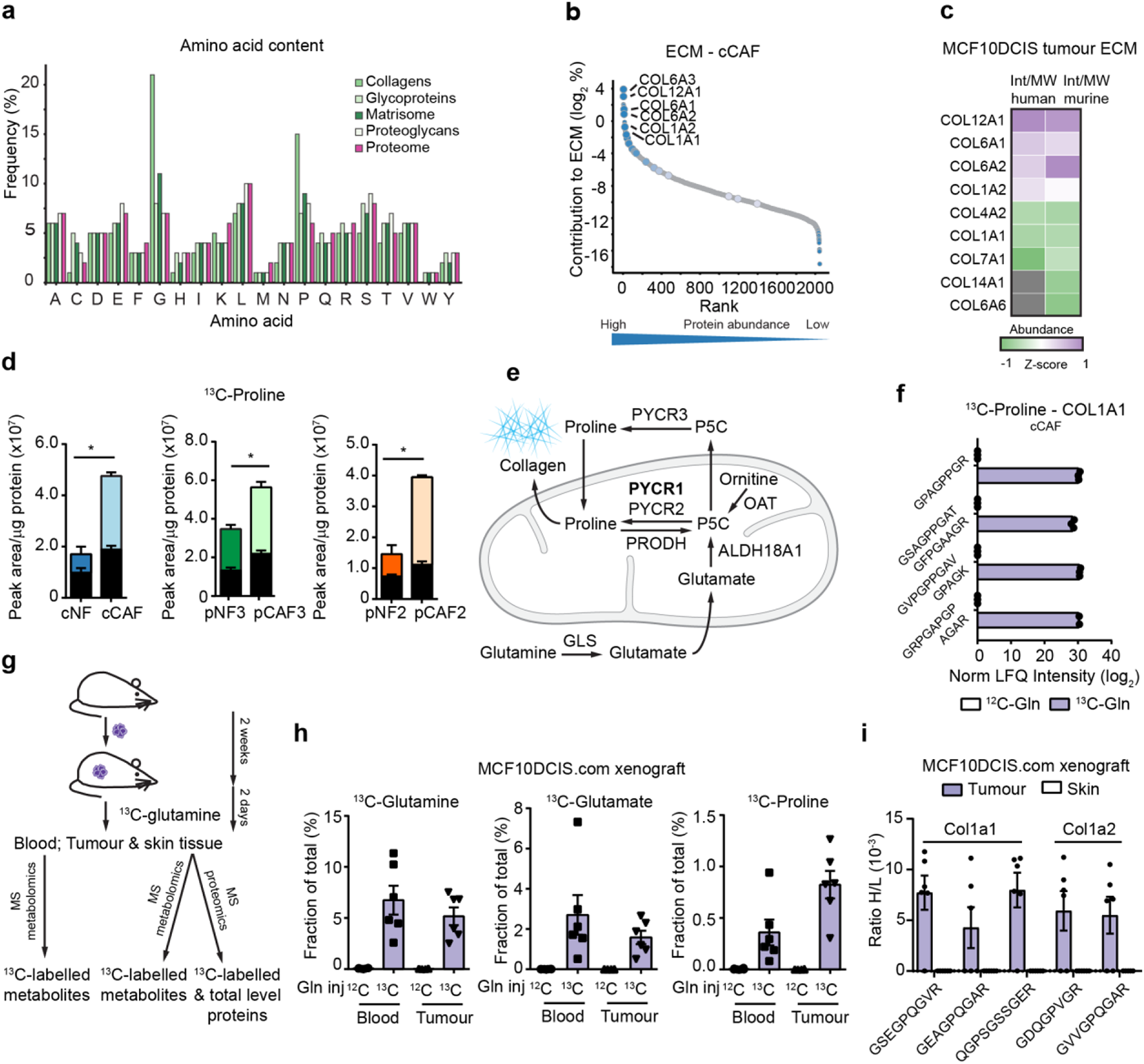
CAFs use glutamine-derived proline for collagen synthesis. **a.** Average frequency of occurrence of each amino acid in proteins in the human proteome (purple), matrisome (dark green) and matrisome components (other greens) as defined by Naba et al. ^60^. **b.** Estimated total abundance (intensity/molecular weight used to rank proteins from low to high abundant, x axis) of proteins identified in the cCAF ECM as measured by MS-proteomics ^22^. On the y axis is the contribution of each protein to the total mass of the ECM. Highlighted are the most abundant proteins collagens. **c.** Comparison of the abundance (intensity/molecular weight) of collagen proteins, measured by MS-proteomics, between endogenous (murine) and transplanted (human) stroma in xenograft tumours of MCF10DCIS.com cells co-transplanted with pCAF2 fibroblasts and grown for two weeks in immunocompromised Balb/c mice. Z-scoring of the protein intensities was performed separately for murine and human collagens. **d**. Total ^13^C-labelled (coloured bar) and unlabelled (black bar) proline in mammary NFs and CAFs labelled with ^13^C-Glutamine, measured by MS-metabolomics. N = 3 biological replicates. **e.** Scheme showing proline biosynthesis pathways from glutamine and ornithine. **f.** MS-proteomic analysis of ECM derived from cCAFs labelled for 72h with 2 mM ^13^C-glutamine or ^12^C-glutamine, showing ^13^C-proline incorporation into COL1A1 peptides. **g.** Workflow of the MCFDCIS.com xenografts with ^13^C_5_-glutamine tracing and MS-based tissue analyses. **h.** Proportion of ^13^C-labelled glutamine, glutamate and proline in the blood and tumours of mice with MCF10DCIS.com xenografts after 48h of treatment with ^13^C-glutamine or ^12^C-glutamine. 0 means that no heavy labelled amino acid had been detected at the MS. **i.** MS-proteomic analysis of tumours and skin from (h) showing ^13^C-proline incorporation into murine Col1a1 and Col1a2 peptides. Error bars indicate mean ± SEM. *p ≤0.05, **p ≤0.01, ***p ≤0.001.

To address this question, we used a previously established model of tumour-promoting mammary CAFs and their normal fibroblast parental line ^22, 25, 59^ (referred to as cCAF and cNF, respectively), and two lines of patient-derived CAFs (pCAFs) and their matched normal fibroblasts (pNFs). cCAF were derived from immortalised human normal mammary fibroblasts that had been activated when co-transplanted with human breast cancer cells in subcutaneous tumours ^25^, while we generated pCAFs and pNFs from surgical samples of patients with breast cancer at an advanced stage. Each pCAF and pNF pair was isolated from breast tumour and normal tumour-adjacent tissue of the same patient, based on histopathological criteria. All the fibroblasts expressed the mesenchymal marker vimentin but not the epithelial marker keratin 18 (**Extended Data Figure 1a,b**). Moreover, both cCAFs and pCAFs have a myofibroblast-like phenotype, as they produce abundant collagen and express high levels of αSMA compared to matched NFs^22, 25^ (**Extended Data Figure 1c-e**). Our previous proteomic analysis of SDS-soluble ECM derived from cCAFs in culture ^22^ showed that collagen makes more than 30% of the total ECM, and that COL12A, COL6A and COL1A are the most abundant collagen proteins (**Figure 1b** and **Supplementary Data S2**). To assess whether CAFs produce a similar ECM also in vivo, we performed MS-proteomic analysis of xenografts grown in Balb/c nude mice following subcutaneous co-transplantation of pCAF and MCF10DCIS.com breast cancer cells, a model in which CAFs accelerate tumour growth ^24^ and that we used later on. This analysis showed that COL12A, COL6A and COL1A were the most abundant collagens produced by both endogenous (murine) and transplanted (human) tumour stroma (**Figure 1c** and **Supplementary Data S3** datasheet Intensity Collagens Tum). Thus, CAFs produce abundant amounts of collagen I and collagen VI, both of which may promote breast cancer progression, and our CAF lines are relevant models to study the role of CAF-derived collagen in tumours.

MS metabolomics analysis of intracellular metabolites in mammary CAFs and NFs cultured in basal conditions (DMEM with 25 mM glucose and 2 mM glutamine) pinpointed proline as being consistently more abundant in CAFs (**Extended Data Figure 1f**). MS-based tracing experiments of cells cultured for 24 hours in DMEM where glucose or glutamine had been replaced with uniformly labelled ^13^C_6_-glucose or ^13^C_5_-glutamine showed that CAFs synthesised more proline from glutamine (**Figure 1d** and **Extended Data Figure 1g,h**). Using the cCAF line, we also found that culturing them in DMEM containing physiological levels of glucose (5 mM) and glutamine (0.65 mM) (Physiol DMEM) had minor impact on the total levels of intracellular proline and did not reduce proline synthesis from glutamine (**Extended Data Figure 1i,j**). Similarly, proline synthesis from glutamine was not altered when physiological levels of proline (200 µM ^62^) were added in the medium (**Extended Data Figure 1k**). Hence, CAFs have an enhanced ability to synthesise proline, which is one of the most abundant amino acid residues in collagen proteins.

To determine whether CAFs utilise glutamine-derived proline to make collagen (**Figure 1e**) we cultured them in medium with ^13^C_5_-glutamine and monitored the presence of ^13^C_5_-proline in collagen proteins by MS-proteomic analysis of their SDS soluble ECM. MS analysis detected ^13^C_5_-proline in COL1A1 peptides (**Figure 1f** and **Supplemental Data S4**), demonstrating that CAFs use newly synthesised proline to make collagen, similarly to normal and TGFβ-activated fibroblasts in culture ^63, 64^. To explore whether this was also the case *in vivo*, we traced ^13^C_5_-glutamine in MCF10DCIS.com tumour-bearing Balb/c nude mice (**Figure 1g**). ^13^C_5_-glutamine, ^13^C_5_-glutamate and ^13^C_5_-proline were detected in blood as well as in tumour tissue (**Figure 1h**). While the fractional enrichment of ^13^C-glutamine and ^13^C-glutamate were comparable in blood and tumour, ^13^C_5_-proline was more abundant in tumour than in blood (1% vs less than 0.5%, respectively), demonstrating that the *in situ* synthesis of proline from glutamine contributes to the intratumoral pool of proline. Furthermore, MS-proteomic analysis of the tumours detected murine Col1a1 and Col1a2 peptides containing ^13^C_5_-proline (**Figure 1i** and **Supplementary data S3** datasheet H/L Ratio Collagen peptides). Together these data provide a first evidence that circulating glutamine is used to supply proline for tumour collagen synthesis *in vivo*. Interestingly, we could not detect ^13^C_5_-proline-containing Col1a1 and Col1a2 peptides in the skin (**Figure 1i**), even though the total levels of collagen were similar to those in tumours (Extended Data Figure 1l and **Supplementary data S3**, datasheets LFQ Intensity Collagens Tum-Skin and H/L Ratio Collagen peptides). This is in agreement with low rate of proline synthesis from glutamine observed in most tissue in non-tumour-bearing mice ^65^ or suggests higher rate of collagen synthesis in tumours than in normal skin. Hence, increased proline synthesis from glutamine is an acquired trait of CAFs, and may support collagen production in tumours.

### PYCR1 is expressed in collagen-producing CAFs and upregulated in aggressive breast cancer stroma

To identify enzymes of the proline biosynthetic pathway responsible for the increased proline synthesis in CAFs (**Figure 1e**), we measured the total proteome of cCAF and cNF (**Supplementary Data S5**). PYCR1-3 enzymes catalyse the last step in proline biosynthesis, by converting pyrroline-5-carboxylate (P5C) into proline. While PYCR1 and PYCR2 are mitochondrial proteins, PYCR3 resides in the cytosol. PYCR1 was the most upregulated enzyme in cCAF (**Figure 2a**). PYCR2 and ALDH18A1, which generates P5C from glutamate, were also upregulated, but to a lower extent (**Figure 2a**), while the levels of PYCR3 and ornithine aminotransferase (OAT), which is involved in the synthesis of proline from urea cycle (**Figure 1e**), did not differ between cCAF and cNF (**Figure 2a**). Western blot analysis confirmed PYCR1 upregulation in all our CAFs compared to their matched NF (**Figure 2b**). Gene expression analysis of stroma microdissected from normal breast and tumour from patients with advanced triple negative breast cancers (TNBC) ^66^ showed that, as in the CAFs, *PYCR1* was the most upregulated enzyme of the proline synthesis pathway in the tumour stroma (**Figure 2c**). Moreover, available gene expression data of patient-matched microdissected stroma and epithelium from normal breast, ductal carcinoma in situ (DCIS) and invasive ductal carcinoma (IDC) ^67^ showed that *PYCR1* expression was significantly higher in DCIS and further increased in IDC, both in the stroma and epithelium (**Figure 2d**). Also *COL1A1* expression significantly increased with disease progression (**Figure 2d**), and positively correlated with *PYCR1* expression in the stroma but not in the epithelium (**Figure 2e**). To determine which cell types contribute to stromal PYCR1 expression, we re-analysed a publicly available single cell RNA sequencing (scRNAseq) dataset of TNBC by Wu et al. ^11^. Among non-epithelial cells, *PYCR1* was expressed in plasma cells and CAFs, particularly a subset of myCAFs, as defined by Wu et al., which express higher levels of *COL1A1* (**Figure 2f,g Extended Data Figure 2a,b**). Conversely, *PYCR2* was found expressed only by few of those CAFs (**Extended Figure 2c**) and *PYCR3* was not detected, in line with our *in vitro* data (**Figure 2a**). *PYCR1* was also expressed in epithelial cells (**Figure 2g**), as previously reported, thus showing that both epithelium and CAFs contribute to *PYCR1* expression in IDC.

**Figure 2.**
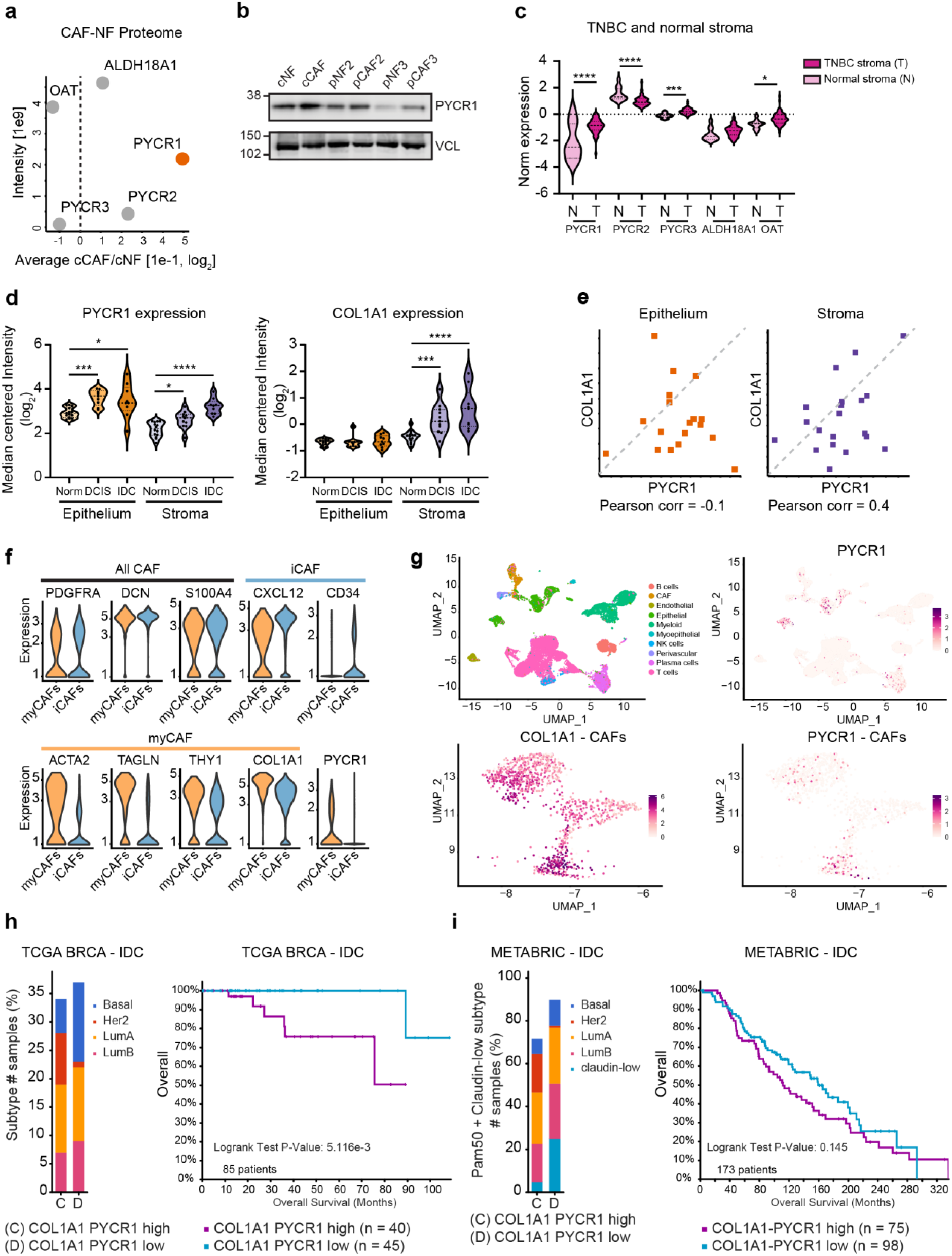
Collagen producing CAFs and stroma of invasive breast cancers express high levels of PYCR1. **a.** Fold change in levels of proline synthesis enzymes between cCAFs and cNFs in an MS-proteomic analysis. In the y axis are reported the Intensities of each protein, as mean of their abundance in the fibroblasts. N = 4 biological replicates **b.** Representative western blot showing PYCR1 levels in paired CAFs and NFs. VCL = vinculin = loading control. **c.** Expression of proline synthesis pathway and stromal markers in LCMD sections of normal breast stroma and TNBC associated stroma from Saleh et al. ^66^. **d.** Violin plots showing the expression levels of *PYCR1* (probe g5902035_3p_a_at) and *COL1A1* (probe Hs.172928.0.A2_3p_a_at) in laser captured microdissected normal breast, ductal carcinoma in situ (DCIS) and invasive ductal carcinoma (IDC) stroma ad epithelium from Ma et al. dataset ^67^. Quantitative data downloaded from Oncomine. Median and quartiles are indicated with dashed lines. Significance was calculated with one-way ANOVA test with Sidak’s multiple comparison test. **e**. Gene expression correlation (Pearson) between PYCR1 and COL1A1 in IDC patients shown in (d). **f.** Violin plots showing the expression of markers commonly associated to myCAF, iCAF and total CAF, and of PYCR1 in triple negative breast cancer (TNBC) tumours from Wu et al. ^11^. **g**. UMAP visualisation of stromal, immune and cancer cells (top plots) aligned using canonical correlation analysis in Seurat. Top left, cells are coloured by their cell type annotation from Wu et al. ^11^. Bottom panels contain only CAFs, as defined by Wu et al. **h, i**. Kaplan-Meier plots comparing overall survival of patients with IDC tumours expressing high or low levels of both *COL1A1* and *PYCR1*. On the left of each curve is shown the distribution of breast cancer subtypes in the two subsets of patients. Data generated with cBioportal using TCGA Pan Cancer Atlas (h) and METABRIC (i).

In light of these results, we explored whether high *PYCR1* and *COL1A1* expression correlated with clinical outcome in breast cancer patients. We analysed a cohort of 780 IDC patients from the Pan Cancer Atlas study of The Cancer Genome Atlas (TCGA) ^68, 69^. Around 10% of the patients expressed either high levels (upper quartile) of both *COL1A1* or *PYCR1* or low levels (lower quartile) of the two genes. The subset with higher levels included a bigger proportion of HER2+ tumours and patients had significantly worse outcome (**Figure 2h**). We obtained similar results analysing IDC patients of the METABRIC study ^70^ (**Figure 2i**). Notably, there was a similar trend of worse outcome when tumours expressed high *PYCR1* and *COL1A1* in other tumour types, including renal, lung, pancreatic, head and neck and colorectal cancer (**Extended Figure 2d**).

Together, these data provide a link between PYCR1 and collagen in CAFs and in breast cancer stroma. Based on these premises, we investigated further the role of PYCR1 in collagen production in CAFs.

### PYCR1 provides proline for collagen biosynthesis

Inhibiting PYCR1 genetically with siRNA or shRNA or pharmacologically with a small molecule inhibitor (PYCR1i) ^71^ was sufficient to decrease newly synthesised ^13^C_5_-proline from ^13^C_5_-glutamine (**Figure 3a-e** and **Extended Data Figure 3a-e**), while minimally affecting CAF proliferation (**Extended Data Figure 3f,g**). Similarly, inhibiting glutaminase (GLS, **Figure 1e**) with the clinical compound CB-839^72^ abrogated synthesis of new proline and collagen in cCAF (**Extended Data Figure 3h,i**). Strikingly, reducing PYCR1 levels in cCAF or pCAF as well as its pharmacological inhibition was sufficient to decrease collagen deposition in the ECM, as measured with the mCherry-tagged collagen probe CNA35 ^73^ and western blot for COL6A1, which was almost fully rescued by providing CAFs with exogenous proline (**Figure 3 f-k** and **Extended Data Figure 3 j,k**). Notably, supplementing the culture medium with supraphysiological concentration of proline ^62^ was able to abrogate the effect of PYCR1 inhibition, while physiological levels had only a minor impact. Addition of exogenous proline did not however increase collagen production in CAFs with no PYCR1 inhibition (**Extended Data Figure 3l)**, suggesting that endogenous PYCR1 activity produces sufficient proline for collagen production in CAFs. Thus, so far, our data show that proline derived from PYCR1 is required for collagen production, particularly when there is limited or physiological availability of exogenous proline. Further supporting this hypothesis, the production of collagen proteins decreased upon silencing PYCR1, while *COL1A1* mRNA levels were not modulated in pCAF and cCAF (**Figure 3l** and **Extended Data Figure 3m**). Moreover, differential ribosome codon reading (diricore) analysis ^74^ showed that reduced levels of PYCR1 in CAFs induced ribosome stalling specifically at proline codons, which was rescued with the addition of exogenous proline (**Extended Data Figure 3n**). Among the mRNAs that were identified and found to be affected by proline levels, there were several collagens, including *COL1A1* (**Figure 3m**). Thus, a major function of PYCR1 in CAFs is to provide proline residues to maintain enhanced production of collagen proteins.

**Figure 3.**
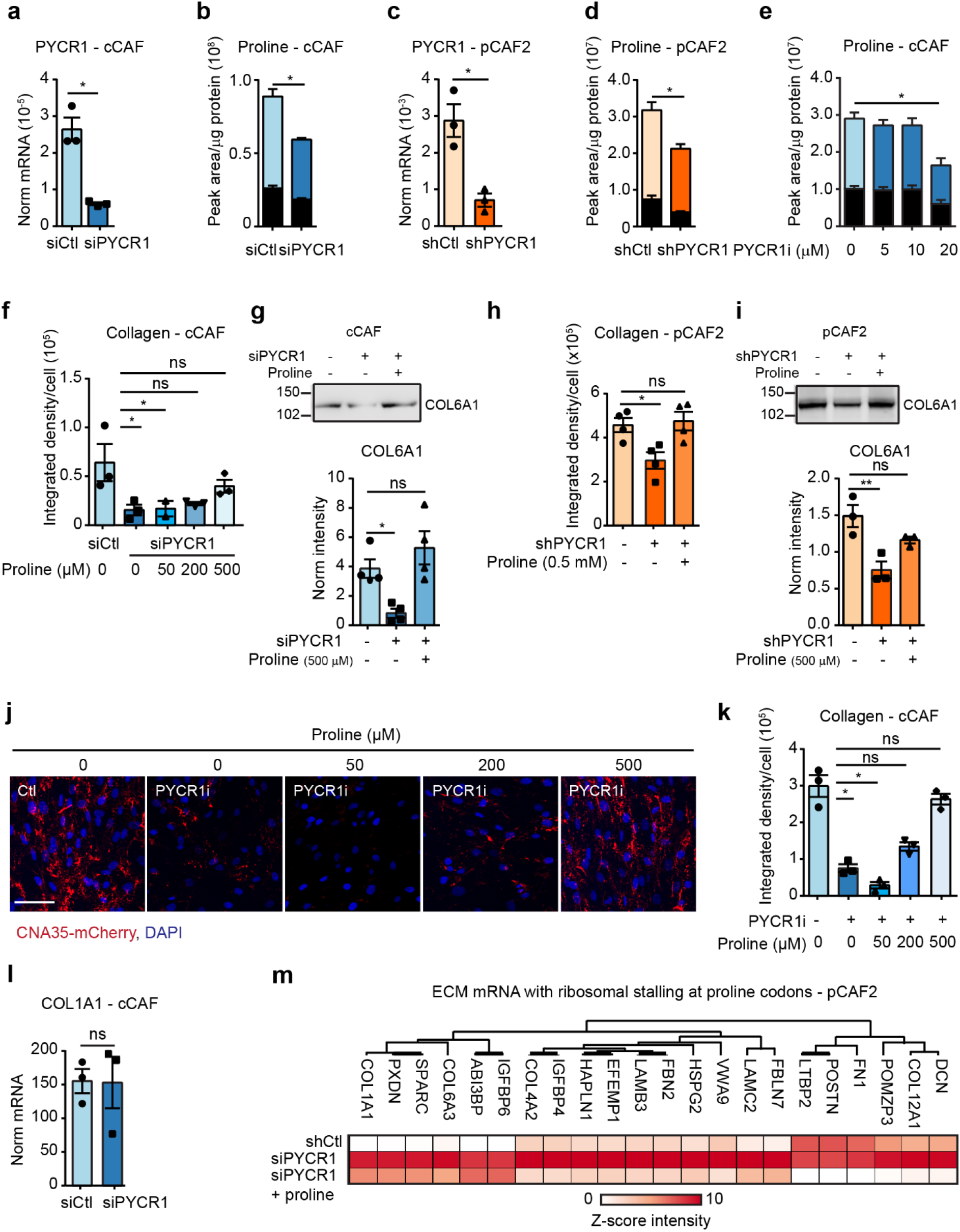
CAFs increase proline biosynthesis via PYCR1. **a.** *PYCR1* expression in cCAF transfected with siCtl or siPYCR1, measured by RT-qPCR and normalised to 18S levels. N = 3 biological replicates. **b.** Total ^13^C-labelled (coloured bar) and unlabelled (black bar) proline measured by MS in cCAFs transfected with siCtl or siPYCR1 and labelled with ^13^C-glutamine. N = 3 biological replicates. **c.** *PYCR1* expression in pCAF2 transfected with shCtl or shPYCR1, measured by RT-qPCR and normalised to 18S levels. N = 3 biological replicates. **d.** Total ^13^C-labelled (coloured bar) and unlabelled (black bar) proline measured by MS in shCtl and shPYCR1 pCAF2 and labelled with ^13^C-glutamine. N = 3 biological replicates **e.** Total ^13^C-labelled (coloured bar) and unlabelled (black bar) proline measured by MS in cCAFs treated with PYCR1i or DMSO, as control, and labelled with ^13^C-glutamine. N = 3 biological replicates. **f, h.** Quantification of CNA35-mCherry labelled collagen produced by cCAFs (f) and pCAF (h) transfected with sh/siCtl or sh/siPYCR1 and treated with proline or PBS as control. N = 3 biological replicates. **g, i**. Representative Western blot and quantification for COL6A1 of decellularised ECM generated from cCAF (g) and pCAF (i) transfected with sh/siCtl or sh/siPYCR1 and treated with proline or PBS as control. COL6A1 signal was normalised by total protein loading measured with Red Ponceau (see Extended Data Figure 9). **j, k.** Representative images (j) and quantification (k) of CNA35-mCherry stained collagen produced by CAFs treated with PYCR1i or DMSO as control and proline or PBS as control. N = 3 biological replicates. **l.** *COL1A1* expression in siCtl and siPYCR1 CAFs, measured by qPCR and normalised to 18S expression. **m.** Diricore analysis of ribosome stalling on proline codons in mRNA of pCAF2 transfected with siCtl or siPYCR1 ± proline, showing ECM mRNAs most affected by PYCR1 knockdown. N = 3 biological replicates. Scale bar = 50 µm. Error bars indicate mean ± SEM. *p ≤0.05, **p ≤0.01.

### Targeting PYCR1 in CAFs reduces tumour growth and metastasis

As collagen influences tumour development, we asked whether targeting PYCR1 in co-culture systems could affect the cancer cells, and co-cultured cCAF with a primary line of breast cancer cells either in a 3D environment, as spheroids, in microfluidic devices in the absence of exogenous matrix ^75^ or in 2D. Imaging analysis of both co-cultures showed that the majority of the collagen co-localised with the GFP-expressing CAFs, indicating that, similarly to *in vivo* ^76^, CAFs are the major source of collagen, while the contribution of the cancer cells is marginal (**Figure 4a,b**). In the spheroid co-culture, collagen production was reduced upon pharmacological inhibition of PYCR1 in a dose-dependent manner, and exogenous proline added to the medium antagonised this effect (**Figures 4c,d**). Similarly, inhibiting PYCR1 in 2D co-culture strongly reduced collagen production, which was rescued with the addition of exogenous proline (**Figure 4e,f** and **Extended Data Figure 4a,b**). Notably, PYCR1 inhibition in the co-culture significantly reduced the proliferation of the cancer cells but not of the CAFs (**Figure 4g,h**), conversely, inhibiting PYCR1 in cancer cells in mono-culture only modestly reduced their proliferation (**Extended Data Figure 4c**). Cancer cells also proliferated less when cultured on the ECM derived from CAFs treated with PYCR1i (**Figure 4i**). Hence, we have revealed a potentially important role of PYCR1 in CAFs to support cancer cell growth via supporting enhanced collagen production.

**Figure 4.**
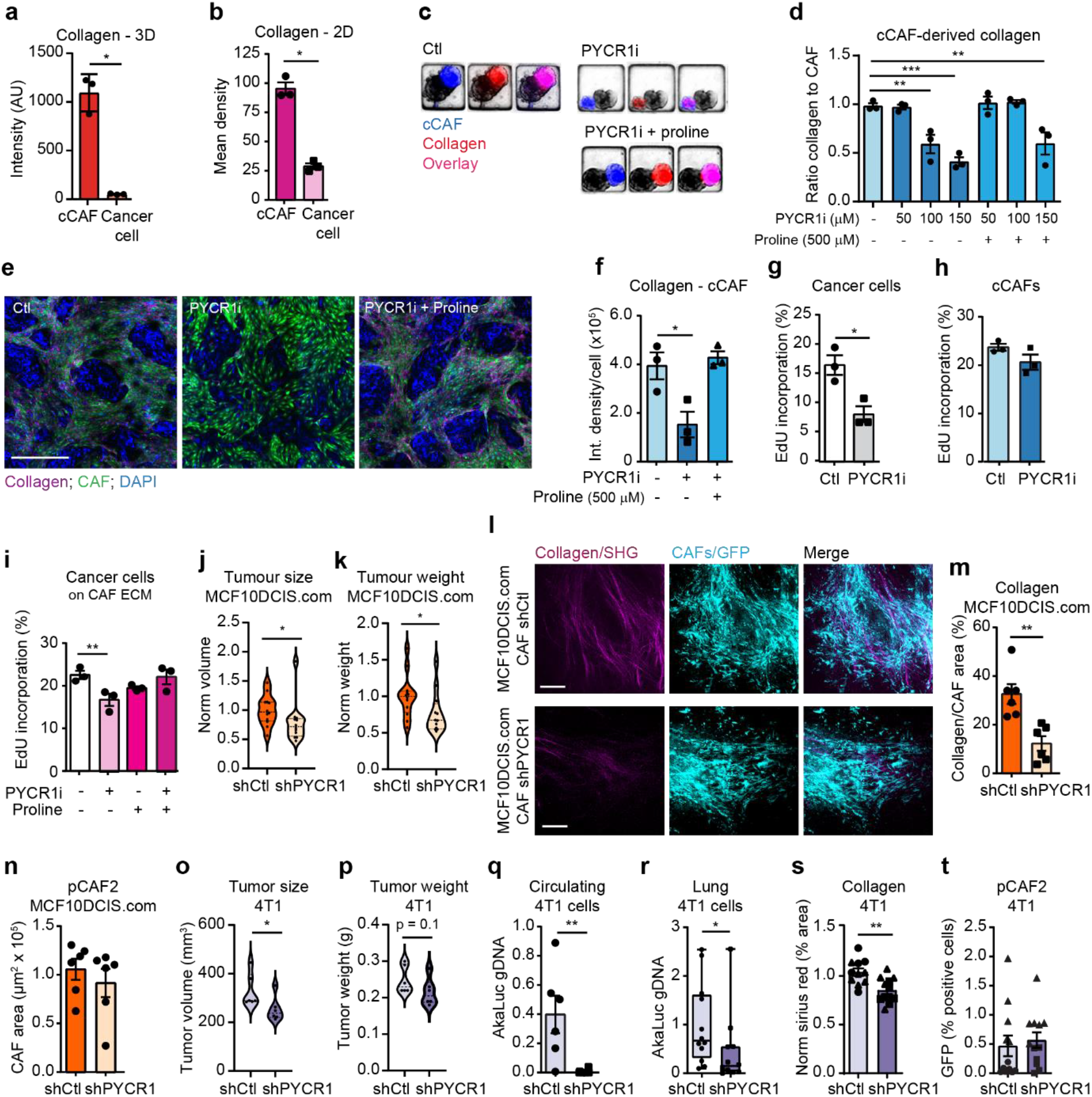
Stromal PYCR1 regulates collagen production and tumour progression *in vivo*. **a, b.** Quantification of collagen produced by cCAFs and breast cancer cells in 3D co-culture (a) and 2D co-culture (b). N = 3 biological replicates. **c, d.** Representative images (c) and quantification (d) of cCAF derived collagen in 3D co-cultures of CAFs (GFP positive) and breast cancer cells treated with PYCR1i or DMSO, as control, and proline. N = 3 biological replicates. **e, f.** Representative images (e) and quantification (f) of cCAF derived collagen in 2D co-cultures of cCAFs (GFP-positive) and cancer cells treated with PYCR1i or DMSO, as control, and proline or PBS, as control. N = 3 biological replicates. **g, h.** EdU incorporation of breast cancer cells (g) and cCAFs (h) in 2D co-cultures treated with PYCR1i or DMSO as control. N = 3 biological replicates. **i.** EdU incorporation of breast cancer cells cultured on ECM derived from CAFs treated with PYCR1i or DMSO, as control, and proline or PBS, as control. Control is the same as in Figure 5v and 6n. N = 3 biological replicates. **j, k.** Tumour volume (j) and weight (k) of xenografts of MCF10DCIS.com cells co-transplanted with pCAF2 shCtl or shPYCR1. Tumours were harvested 14 days after injections. N = 12 mice for each condition from two independent experiments (6 mice/experiment). **l, m.** Representative images (l) and quantification (m) by SHG signal of fibrillary collagen surrounding shCtl or shPYCR1 CAFs in MCF10DCIS.com xenografts from (k). N = 6 mice from each condition from one experiment. **n.** Area of shCtl or shPYCR1 CAFs (based on GFP signal) in images used for (m). **o, p.** Tumour volume (o) and weight (p) of tumours from 4T1 cells expressing Aka-luciferase co-transplanted with pCAF2 shCtl or shPYCR1. N = 6 mice for each condition from one experiment. **q.** Quantification of circulating tumour cell DNA by RT-qPCR in blood taken from mice with tumours of 4T1 cells expressing Aka-luciferase (AkaLuc) co-transplanted with pCAF2 shCtl or shPYCR1. Data was normalised to beta-actin (ACTB) levels. N = 6 mice for each condition from one experiment. **r.** Quantification of tumour cell DNA by RT-qPCR in lungs from mice with tumours of 4T1 cells expressing Aka-luciferase (AkaLuc) co-transplanted with pCAF2 shCtl or shPYCR1. Data was normalised to beta-actin (ACTB) levels. N = 12 mice for each condition from two experiments (6 mice/group/experiment). **s.** Quantification of Sirius Red collagen staining in FFPE fixed sections of 4T1 expressing Aka-Luc + CAF shCtl/shPYCR1 tumours. N = 12 mice for each condition from two independent experiments (6 mice/group/experiment). **t.** Quantification (based on IHC staining for GFP) of shCtl or shPYCR1 CAFs in FFPE fixed sections of 4T1 expressing Aka-Luc + CAF shCtl/shPYCR1 tumours. N = 12 mice for each condition from two independent experiments (6 mice/group/experiment). Error bars indicate mean ± SEM. *p ≤0.05, **p ≤0.01, ***p ≤0.001. Scale bar = 50 µm.

Next, we explored the consequences of targeting PYCR1 in CAFs *in vivo*. MCF10DCIS.com cells were co-transplanted with pCAFs expressing normal (shCtl) or reduced levels of PYCR1 (shPYCR1) subcutaneously in Balb/c nude mice, and tumours harvested two weeks after transplantation. While this model does not recapitulate the heterogeneity of the CAFs (i.e. iCAF and myCAF) found in patients, we reasoned it to be a useful model to study the consequences that targeting pathways that support ECM-production in CAFs have on tumours progression. Tumours containing CAFs shPYCR1 had significantly reduced size and weight compared to the ones containing CAFs shCtl (**Figure 4j,k**). Moreover, microscopy analysis of the tumours showed a marked decrease in fibrillar collagen deposited around pCAF shPYCR1 compared with pCAF shCtl, while the tumour area covered by CAF was not affected (**Figure 4l-n**), consistent with our *in vitro* data (**Figure 4c-f**). Next, to assess whether targeting PYCR1 in CAFs had an impact on metastasis, we co-transplanted pCAFs shCtl or shPYCR1 with metastatic 4T1 breast cancer cells expressing aka-luciferase in NRMI nu/nu mice, and sacrificed mice twelve days after transplantation. Similarly to the MCF10DCIS.com xenograft model, the size and weight of the tumours containing CAFs shPYCR1 was reduced, although the weight did not reach statistical significance (**Figure 4o,p**). Notably, the co-transplantation of pCAF shPYCR1 significantly reduced the presence of circulating cancer cells in the blood (**Figure 4q**), as well as the presence of metastatic cancer cells in the lungs (**Figure 4r**). Tumours grown with pCAFs shPYCR1 contained less collagen than those grown with pCAFs shCtl (**Figure 4s** and **Extended Data Figure 4d**), while the amount of GFP-expressing CAFs was similar (**Figure 4t**). Conversely, the total amount of blood vessels (Pecam1 staining) and hypoxia (pimonidazole staining) were similar in all the tumours (**Extended Data Figure 4e,f**). These data suggest that PYCR1 in the stroma may control metastatic cancer cell intravasation in the tumour blood vessels, at least in part, via regulating collagen deposition in the tumour stroma.

Thus, our data show that PYCR1 in mammary CAFs represents a stromal vulnerability for ECM production and can be targeted to reduce tumour growth and metastasis.

### CAFs have hyperacetylated histone 3

We next sought to determine the signalling that supports increased collagen production and PYCR1 expression in CAFs. Our MS-metabolomic analyses showed that the total levels of acetyl-CoA were consistently higher in CAFs than in NFs (**Extended Data Figure 1f** and **Figure 5a**). Acetyl-CoA is a signalling molecule and acetyl donor for acetylation of proteins and histones, and is therefore a central epigenetic regulator ^77, 78^. Intriguingly, MS-based global acetylation analysis of cCAF and cNF showed histone H3 more acetylated in CAFs at sites K18, K23 and K27 (**Figure 5b** and **Supplemental Data S5**), which are substrates of the epigenetic regulator histone acetyl-transferase EP300 ^79^. We confirmed that H3K27 was hyperacetylated in all our CAF lines, when compared to their matched NF, by western blot analysis (**Figure 5c**). We focused on H3K27 because its hyperacetylation is an established marker of enhancer activity ^80^, and increased levels of nucleo-cytosolic acetyl-CoA has been shown to promote its acetylation through the activation of EP300 ^81, 82^. Notably, the recruitment of EP300 and hyperacetylation of histone H3 at enhancers of pro-fibrotic genes, including collagen, are a landmark event in fibrosis, and are involved in TGFβ-regulated gene expression, which is a major trigger of fibroblast activation ^83–85^. Furthermore, there is increasing evidence that an epigenetic switch underpins CAF activation ^54, 57, 86^, which would explain why CAFs maintain acquired functions, such as increased ECM production (**Extended Data Figure 1d,e**), despite long-term culture *in vitro* without stimulation from cancer cells ^22, 23, 87^. Based on these grounds, we sought to investigate the link between EP300, acetyl-CoA levels and histone acetylation with collagen production in CAFs.

**Figure 5.**
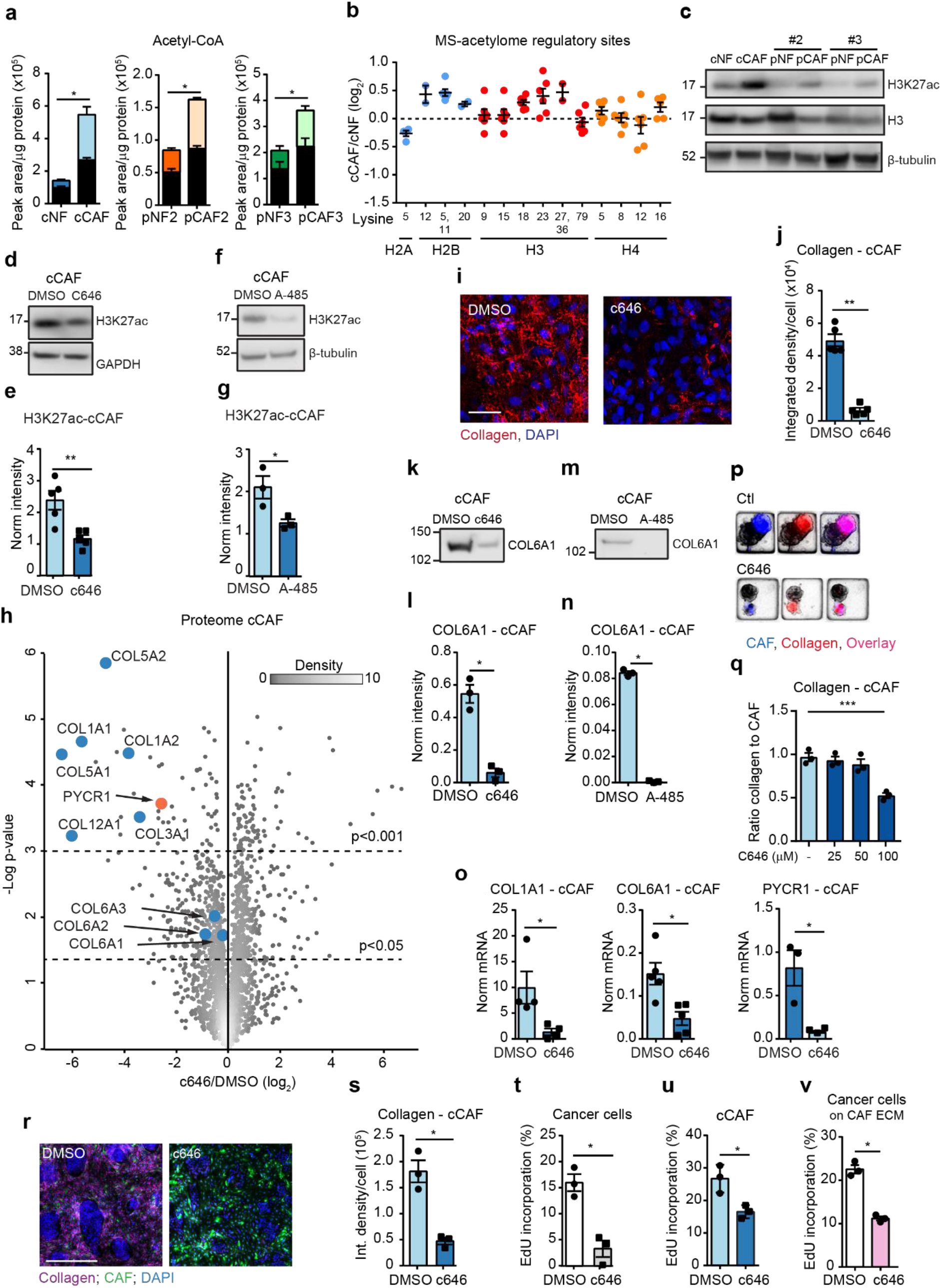
H3 acetylation regulates collagen production in CAFs. **a.** ^13^C-labelled (coloured bar) and unlabelled (black bar) acetyl-CoA measured by MS in cCAFs and NFs labelled with ^13^C-glucose. N = 3 biological replicates. **b.** SILAC ratios of histone acetylation sites with a known regulatory function (based on PhosphoSitePlus, release 2016) identified in the MS-based acetylome of cNFs and cCAFs. N = 5 biological replicates. **c.** Representative western blot showing H3K27ac levels in mammary NFs and CAFs. Total H3 and β-tubulin were used as loading controls. **d, e.** Representative western blot (d) and quantification (e) showing H3K27ac levels in cCAFs treated with c646 or DMSO as control. GAPDH was used as loading control. N = 5 biological replicates. **f, g.** Representative western blot (f) and quantification (g) showing H3K27ac levels in cCAFs treated with A-485 or DMSO as control for 48 h. β-tubulin was used as loading control. N = 3 biological replicates. **h.** Volcano plot showing the average log_2_ ratios of proteins quantified in the total proteome of cCAFs treated with c646 or DMSO as control. N = 3 biological replicates. **i, j.** Representative images (i) and quantification (j) of collagen produced by cCAFs treated with c646 or DMSO as control. N = 5 biological replicates. **k, l.** Representative western blot (k) and quantification (l) of COL6A1 in decellularised ECM derived from cCAFs treated with c646 or DMSO as control. N = 3 biological replicates. **m, n.** Representative western blot (m) and quantification (n) of COL6A1 in decellularised ECM derived from cCAFs treated with A-485 or DMSO as control. N = 3 biological replicates. **o.** mRNA expression of *COL1A1*, *COL6A1* and *PYCR1* in cCAFs treated with c646 or DMSO as control. N ≥ 3 biological replicates. **p, q.** Representative images (p) and quantification (q) of collagen in 3D spheroid co-cultures of CAFs (GFP positive) and breast cancer cells treated with c646 or DMSO as control. N = 3 biological replicates. **r, s.** Representative images (r) and quantification (s) of collagen in 2D co-cultures of CAFs (GFP positive) with breast cancer cells with c646 treatment or DMSO as control. N = 3 biological replicates. **t.** EdU incorporation by breast cancer cells in 2D co-culture with CAFs treated with c646 or DMSO as control. N = 3 biological replicates. **u.** EdU incorporation by CAFs in 2D co-culture with breast cancer cells treated with c646 or DMSO as control. N = 3 biological replicates. **v.** EdU incorporation by breast cancer cells seeded on ECM derived from CAFs that were treated with c646 or DMSO as control. Control is the same as in Figure 4i and 6n. N = 3 biological replicates. N = 3 biological replicates. Collagen was visualised with the collagen binding protein CNA35-mCherry. Error bars indicate mean ± SEM. *p ≤0.05, **p ≤0.01, ***p ≤0.001. Scale bar = 200 µm. See Extended Data Figure 9 for red Ponceau staining of the blots used for COL6A1 staining in the ECM, which was used to normalise to total protein content in each lane.

### EP300 supports collagen production in CAFs

First, we assessed the role of EP300 using the cCAF line. Treating CAFs with the EP300 inhibitors c646 ^81, 88^ or A-485 ^89^ significantly reduced H3K27 acetylation (**Figure 5d-g** and **Extended Data Figure 5a**). To determine whether this reduction in histone acetylation corresponded to a regulation in collagen production, we performed a MS-proteomic analysis of cCAFs after 24h treatment with c646 (**Supplementary Data S6**). Strikingly, collagen proteins, including COL1A and COL6A, were among the most highly downregulated by c646 treatment, and PYCR1 was also strongly downregulated (**Figure 5h**). We further confirmed reduced collagen deposition in the ECM upon EP300 inhibition by microscopy (**Figure 5i,j**) and western blot (**Figure 5k-n**). Moreover, inhibiting EP300 decreased expression of both collagens and *PYCR1* at the transcriptional level (**Figure 5o**), suggesting that EP300 is an epigenetic regulator of collagen production in CAFs. The fact that *PYCR1* and collagen levels were co-regulated further supports our evidence that proline synthesis is upregulated to support collagen production in CAFs.

CAFs also deposited less collagen when treated with c646 in 3D co-culture with cancer cells, and this response was dose-dependent (**Figure 5p,q**). Similarly in 2D co-culture, inhibiting EP300 strongly inhibited collagen deposition (**Figure 5r,s**) and significantly reduced cancer cell and CAF proliferation (**Figure 5t,u**). Inhibiting EP300 also completely blocked cancer cell proliferation when in mono-culture (**Extended Data Figure 4c**), in line with previous reports ^88^. However, cancer cell proliferation was also inhibited when cells were cultured on ECM derived from CAFs treated with c646 (**Figure 5v**). Therefore, while the reduction in collagen production may have contributed to the proliferation phenotype of the cancer cells reported in the co-cultures, it is likely that the overall phenotype has been determined also by a direct effect of the drug on the cancer cells. Interestingly, cancer cell proliferation was less affected upon treatment in the co-culture than in the mono-culture (**Extended Data Figure 4c** and **Figure 5t**), suggesting that CAFs may have a protective effect against c646 treatment. Overall, here we show that EP300 activity is a major regulator of collagen production in CAFs that can be targeted to reduce cancer cell growth.

### Nucleo-cytosolic acetyl-CoA levels regulate collagen production in CAFs

Since EP300 activity is dependent on nucleo-cytosolic acetyl-CoA levels ^81, 82^, we determined the source of acetyl-coA in the CAFs. Tracing experiments of cCAFs exposed for 24 hours to uniformly heavy carbon-labelled metabolites showed that glucose contributed around 50% of the total acetyl-CoA, while acetate, glutamine and pyruvate contributed to a much lesser extent (each ∼ 10% of total acetyl-CoA), with minor differences depending on whether cells were cultured in DMEM or Physiol DMEM, which contains physiological levels of glucose (5 mM), glutamine (0.65 mM), acetate (100 µM) and pyruvate (100 µM) (**Figure 6a** and **Figure 5a**). Hence, we conclude that in cultured CAFs most of the acetyl-CoA is synthesised from glucose-derived pyruvate via the pyruvate dehydrogenase complex (PDC) ^90, 91^ (**Figure 6b**). There is evidence that PDC can translocate into the nucleus to synthesise acetyl-CoA *in loco* ^92, 93^, but we excluded that this occurs in CAFs because we could not detect PDHA1 in the mitochondria, by immunofluorescence staining of fixed cells and western blot analysis of their mitochondria (**Extended Data Figure 5b,c**). The efflux of acetyl-CoA from the mitochondria to the cytosol occurs in the form of citrate via the citrate carrier SLC25A1 and is converted back to acetyl-CoA by the of ATP citrate synthase (ACLY) either in the cytosol or in the nucleus (**Figure 6b**). In cCAF, ACLY was not detected in the nucleus by subcellular fractionation (**Extended Data Figure 5d**), suggesting that acetyl-CoA is primarily made in the cytosol. However, small molecules such as acetyl-coA can freely enter the nucleus through nuclear pores ^94^. To assess whether the levels of nucleo-cytosolic acetyl-CoA influence H3K27 acetylation in CAFs, we used a previously established model, in which nucleo-cytosolic acetyl-CoA is reduced by inhibiting ACLY with the inhibitor BMS303141, and then replenished by adding exogenous acetate at supra-physiological levels (1mM). Acetate is then converted to acetyl-CoA by the acetyl-coenzyme A synthetase (ACSS2) ^81, 82^ (**Figure 6b**). BMS303141 treatment for 24 hours at 50 µM effectively inhibited ACLY, as it induced accumulation of ^13^C_2_-citrate derived from ^13^C_6_-glucose, while reducing levels of ^13^C_2_-labelled and total acetyl-CoA, and total acetyl-CoA levels were restored with acetate supplementation (**Extended Data Figure 5e-g**). Inhibiting ACLY consistently reduced H3K27 acetylation, which was recovered by adding acetate to the culture medium (**Figure 6c,d**). Notably, stimulating untreated CAFs with supra-physiological levels of acetate did not increase histone acetylation (**Extended data Figure 5h,i**), suggesting that while acetate/ACSS2 promote H3K27 acetylation when other sources of acetyl-CoA are limited, they are not a driving factor in the increase in histone acetylation in CAFs. Inhibiting ACLY in cCAF (**Figure 6e**) and pCAF (**Extended Figure 5j**) additionally reduced mRNA expression of both *PYCR1*, *COL1A1* and *COL6A1*, and PYCR1 protein levels (**Extended data Figure 5k**), which were rescued with acetate, supporting that acetyl-coA levels control collagen production at epigenetic level. BMS303141 treatment also reduced collagen production in a dose-dependent manner when CAFs were co-cultured with breast cancer cells in 3D (**Figure 6f,g**) and in 2D (**Figure 6h-k**). Stimulating CAFs with acetate alone had no effect on collagen production (**Extended data Figure 5l**), corresponding with the lack of effect on H3K27 acetylation (**Extended Data Figure 5h,i**). Similarly to inhibiting collagen production by targeting PYCR1, BMS303141 opposed cancer cell proliferation, and acetate abrogated this effect, while it had no significant impact on CAF growth (**Figure 6l,m**). Of note, BMS303141 treatment of cancer cells cultured alone had minor impact on their proliferation (**Extended data Figure 4c**). Moreover, cancer cells grew significantly less on ECM derived from BMS303141-treated CAFs (**Figure 6n**). Thus, nucleo-cytosolic levels of acetyl-CoA in CAFs influence production of collagen, which may support cancer cell growth, and can be effectively inhibited by targeting ACLY.

**Figure 6.**
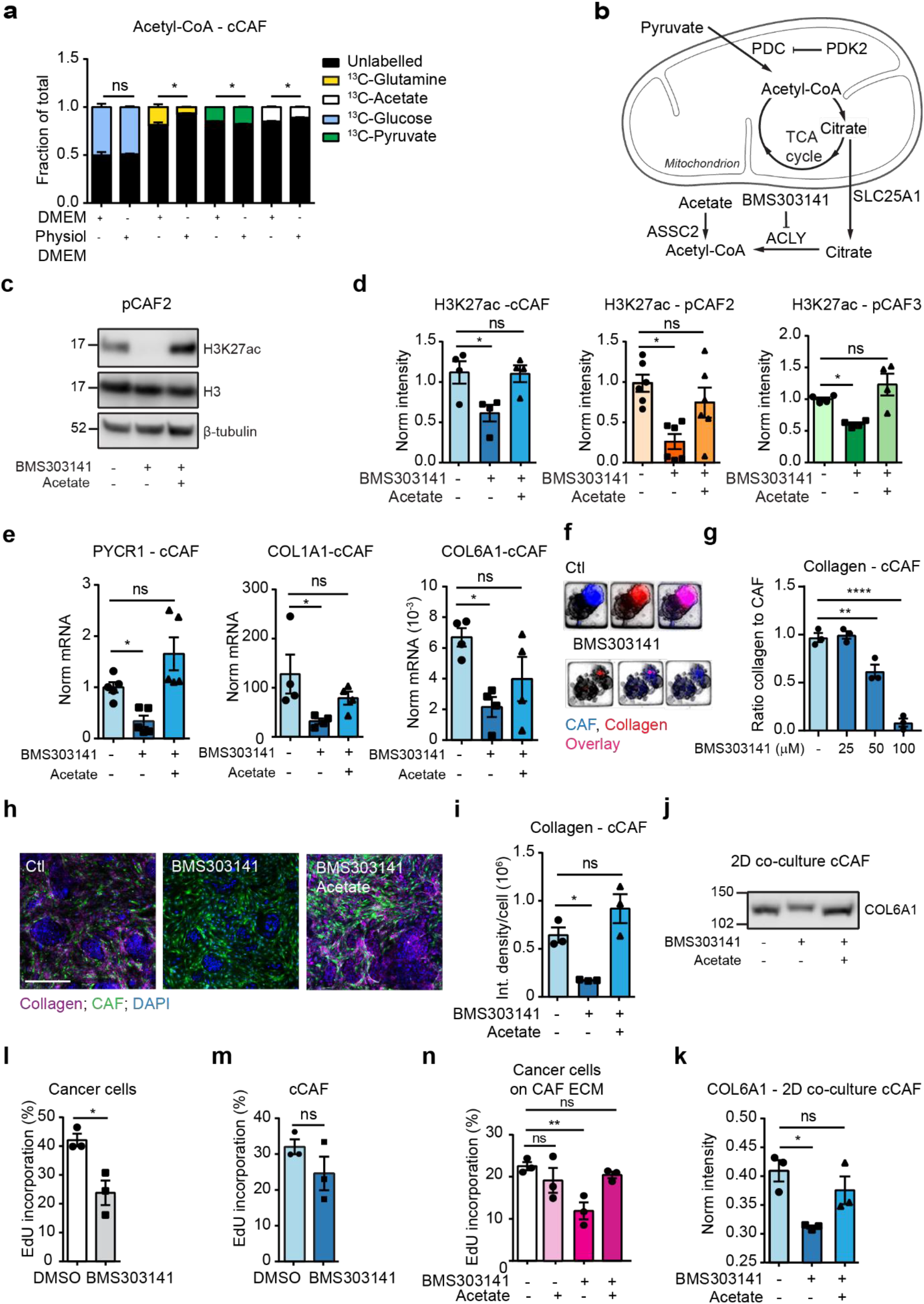
Nucleo-cytosolic acetyl-coA regulates collagen production. **a.** Incorporation of ^13^C-labelled glucose, glutamine, pyruvate or acetate into acetyl-coA in mammary cCAFs cultured in standard DMEM (25 mM glucose, 2 mM glutamine, 1 mM pyruvate, 100 µM acetate) or physiological DMEM (5 mM glucose, 0.65 mM glutamine, 100 µM pyruvate, 100 µM acetate). **b.** Scheme showing acetyl-CoA production and export out of mitochondria. **c, d.** Representative western blot (c) and quantification (d) showing H3K27ac levels in mammary CAFs following treatment with DMSO as control or BMS303141 and acetate or PBS, as control. β-tubulin was used as loading control. N = 4-6 biological replicates. **e.** mRNA expression of *PYCR1, COL1A1* and *COL6A1* in cCAFs treated with DMSO, as control, or BMS303141 and acetate or PBS, as control. N = 3 biological replicates. **f, g.** Representative images (f) and quantification (g) of collagen in 3D spheroid co-cultures of cCAFs and breast cancer cells treated with DMSO, as control, or BMS303141 and acetate or PBS, as control. N = 3 biological replicates. **h, i.** Representative images (h) and quantification (i) of collagen in 2D co-cultures of cCAFs (GFP positive) with breast cancer cells treated with DMSO, as control, or BMS303141 and acetate or PBS, as control. N = 3 biological replicates. **j, k.** Representative western blot (j) and quantification (k) of COL6A1 in ECM derived from cCAFs following treatment with DMSO, as control, or BMS303141 and acetate or PBS, as control. N = 3 biological replicates. **l.** EdU incorporation by breast cancer cells in 2D co-culture with cCAFs (GFP positive) treated with DMSO as control or BMS303141. N = 3 biological replicates. **m.** EdU incorporation by cCAFs in 2D co-culture with cancer cells ± BMS303141 treatment. N = 3 biological replicates. **n.** EdU incorporation by breast cancer cells seeded on decellularised ECM derived from cCAFs treated with DMSO, as control or BMS303141 and acetate or PBS, as control. Control is the same as in Figure 4i and 5v. N = 3 biological replicates. Collagen was visualised with the collagen binding protein CNA35-mCherry. Error bars indicate mean ± SEM. *p ≤0.05, **p ≤0.01, ***p ≤0.001. Scale bar = 200. See Extended Data Figure 9 for red Ponceau staining of the blots used for COL6A1 staining in the ECM, which was used to normalise to total protein content in each lane.

### PDK2 is a key regulator of acetyl-CoA levels in CAFs

Our MS metabolomic data indicated that PDC activity is the major source of increased acetyl-CoA in CAFs (**Figure 5a** and **Figure 6a**). PDC activity is negatively regulated by pyruvate dehydrogenase kinase proteins (PDK 1-4), which phosphorylate the pyruvate dehydrogenase E1 component subunit alpha A1 (PDHA1) ^90, 91^. Intriguingly, computational analysis of our unbiased phosphoproteomic analysis of cCAF and cNF in basal culture conditions (**Supplemental Data S7**) predicted PDK2 to be the most de-activated kinase in cCAF (**Supplemental Data S8** and **Figure 7a**). Concordantly, the phosphorylation level of PDHA1 at the regulatory site serine 293 was significantly lower in CAFs than in their NF counterparts (**Figure 7b,c** and **Supplemental Data S7**), PDC activity was higher in CAFs (**Figure 7d**) and CAFs had lower levels of PDK2 than their matched NF counterpart (**Figure 7e**). When we assessed the expression of all four PDKs by qPCR, *PDK2* was the most highly expressed in NFs, and strongly downregulated in CAFs (**Figure 7f**). Hence, CAFs express low *PDK2* at both gene and protein levels. CAFs also expressed low levels of *PDK1*, *PDK3* and *PDK4*, and there was no consistent significant difference in expression between CAFs and matched NFs, both at protein and mRNA level (**Figure 7f** and **Extended Data Figure 6a**). Gene expression data of microdissected stroma from normal breast and TNBC tissues ^66^ showed significant downregulation of *PDK2* in the tumour stroma (**Figure 7g**) and that *PDK1*, *PDK3* and *PDK4* were expressed at similar or lower levels than *PDK2* in the tumour stroma, similarly to our CAFs (**Figure 7f**). Re-analysis of available scRNAseq data of TNBC ^11^ further showed that all four PDKs could be detected only in few of the CAFs expressing higher levels of collagen (**Extended Data Figure 6b** and **Extended Data Figure 2b**). Moreover, PDK4 was expressed also by perivascular and endothelial cells, which may explain the high PDK4 levels measured in the stroma of normal breast (**Figure 7g**).

**Figure 7.**
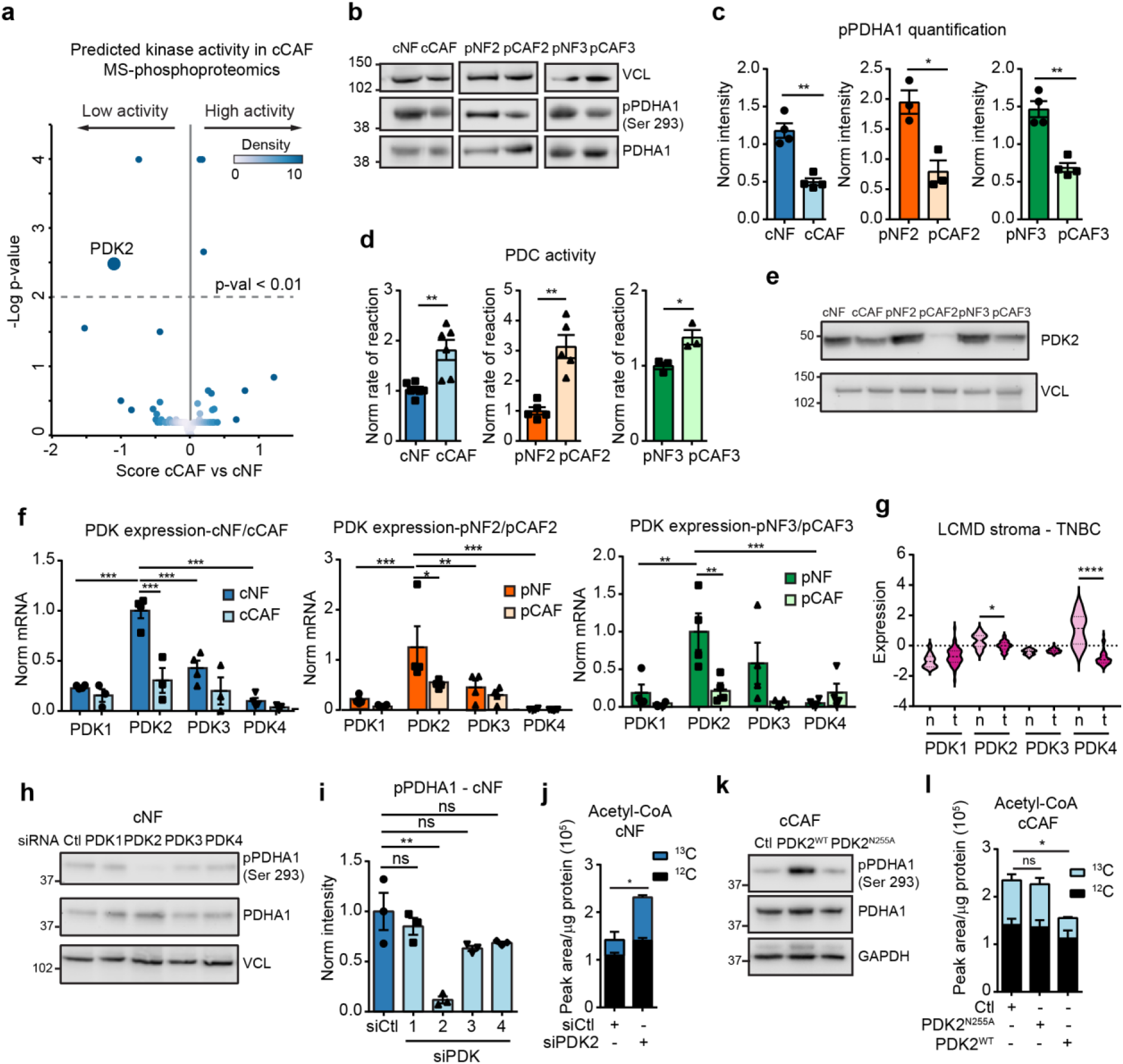
PDK2 regulates PDH activity and acetyl-coA production in CAFs. **a.** Predicted kinase activity in cCAFs compared to cNFs based on the modelling of their MS-based phosphoproteomic data. **b, c.** Representative western blots (b) and quantification (c) showing PDHA1 phosphorylation levels at the regulatory site serine 293 in mammary NFs and CAFs. VCL = Vinculin was used as a loading control. N = 3 or 4 biological replicates. **d.** Pyruvate dehydrogenase activity of NFs and CAFs measured as the rate of NAD+ reduction *in vitro*. N = 3-6 biological replicates. **e.** Representative western blot showing PDK2 levels in paired mammary NFs and CAFs. VCL = Vinculin was used as a loading control. **f.** *PDK1-4* expression in mammary NFs and CAFs in culture, measured by qPCR and normalised to 18S expression. N = 4 biological replicates. **g.** *PDK1-4* mRNA expression in LCMD sections of normal and triple negative breast cancer (TNBC)-associated stroma from Saleh et al.^66^. **h, i.** Representative western blot (h) and quantification (i) showing PDHA1 phosphorylation levels at serine 293 in cNFs transfected with siCtl or siPDK1-4. N = 3 biological replicates. VCL = Vinculin was used as a loading control. **j.** Intracellular acetyl-CoA unlabelled (black) and ^13^C_2_-labelled (coloured) from ^13^C_6_-glucose measured by MS in cNFs transfected with siCtl or siPDK2. N = 3 biological replicates. **k**. Representative western blot showing PDHA1 phosphorylation levels in cCAFs transfected with empty vector, pGC-PDK2^N255A^ or pGC-PDK2^WT^. GAPDH was used as a loading control. **l.** ^13^C_2_-labelled (coloured bar) and unlabelled (black bar) acetyl-CoA measured by MS in cCAFs transfected with empty vector, pGC-PDK2^N255A^ or pGC-PDK2^WT^ and labelled with ^13^C_6_-glucose. N = 3 biological replicates. Error bars indicate mean ± SEM. *p ≤0.05, **p ≤0.01, ***p ≤0.001. Collagen was visualised with the collagen binding protein CNA35-mCherry.

Next, we silenced each of the PDKs in NFs, and while all of them were efficiently silenced (**Extended Data Figure 6c,d**), only silencing PDK2 reduced PDHA1 phosphorylation at its active site and production of PDC-derived ^13^C_2_-acetyl-CoA from ^13^C_6_-glucose both in cNF (**Figure 6h-j**) and pNF (**Extended Data Figure 6e-h**). The lack of effect of PDK1, 3 and 4 silencing on PDHA1 phosphorylation was not due to any compensation by PDK2 expression, as PDK2 protein levels were unaffected by siRNA knockdown of the other PDKs (**Extended data Figure 6i**).

Notably, overexpression of PDK2 wild type (PDK2^WT^), but not a mutant enzymatically inactive form ^95^ (PDK2^N255A^) (**Extended Data Figure 6j,k**), increased PDHA1 phosphorylation, as well as the production of acetyl-CoA via PDC activity in cCAF (**Figure 6k,l**) and pCAF (**Extended Data Figure 6l,m**). Hence, PDK2 is the most expressed of the PDKs in mammary fibroblasts in vitro and a major regulator of PDC activity and acetyl-CoA levels.

### PDC activity promotes enhanced collagen production in fibroblasts

Next, we sought to determine whether the modulation of PDC activity was sufficient to control collagen and proline production. Overexpression of PDK2 in cCAF or pCAF to inactivate PDC significantly reduced H3K27 acetylation (**Figure 8a-d**), as well as expression of *COL1A1*, *COL6A1* and *PYCR1* (**Figure 8e,f**) and collagen deposition in the ECM (**Figure 8g**), and this effect was counteracted by the addition of exogenous acetate (**Figure 8a-g** and **Extended data Figure 7a**). Inhibiting PDC pharmacologically with the clinical compound CPI-613 gave similar results (**Extended Data Figure 7b-f**). Conversely, exogenous acetate did not significantly increase H3K27 acetylation in CAFs expressing inactive PDK2^N255A^ (**Figure 8a-d**). Exogenous acetate was however able to induce H3K27 acetylation, expression of collagen and *PYCR1*, and collagen deposition in cNF (**Extended Figure 7g-k**), showing that increasing acetyl-CoA levels alone is sufficient to induce collagen production in normal fibroblasts. Silencing PDK2 in cNF or pNF to increase PDC activity also enhanced H3K27 acetylation (**Figure 8h-k**), *COL1A1, COL6A1* and *PYCR1* expression (**Figure 8l,m**), and collagen deposition in the ECM (**Figure 8n,o** for cNF and **Extended data figure 7l** for pNF). Notably, these effects were abrogated when inhibiting EP300 with c646 (**Figure 8h-o** and **Extended Data Figure 7l**). Hence, PDC activity is sufficient to support collagen production and *PYCR1* expression and it requires active EP300.

**Figure 8.**
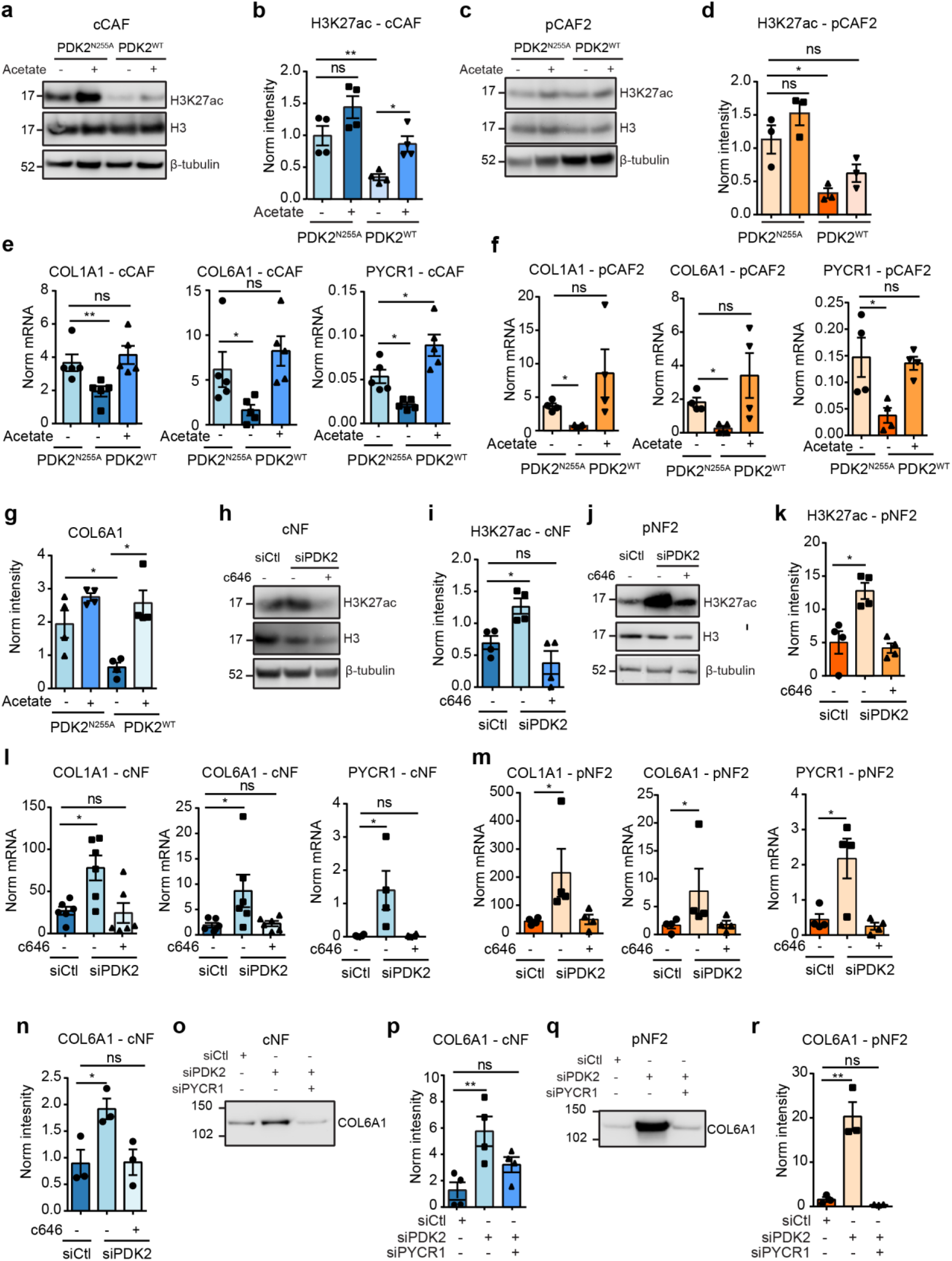
PDH activation regulates collagen production in CAFs. **a, b.** Representative western blot (a) and quantification (b) of H3K27ac levels in cCAFs transfected with pGC-PDK2^N255A^ or pGC-PDK2^WT^ and treated with acetate or PBS as control. N = 4 biological replicates. β-tubulin was used as a loading control. **c, d.** Representative western blot (c) and quantification (d) of H3K27ac levels in pCAF2 transfected with pGC-PDK2^N255A^ or pGC-PDK2^WT^ and acetate or PBS, as control, treatment. N = 3 biological replicates. β-tubulin was used as a loading control. **e.** mRNA expression of *COL1A1*, *COL6A1* and *PYCR1* in cCAFs transfected with pGC-PDK2^N255A^ or pGC-PDK2^WT^ ± acetate treatment. N = 3-5 biological replicates. **f.** mRNA expression of *COL1A1*, *COL6A1* and *PYCR1* in pCAF2 transfected with pGC-PDK2^N255A^ or pGC-PDK2^WT^ and acetate or PBS, as control, treatment. N = 3 biological replicates. **g.** Quantification of COL6A1 levels in ECM from cCAFs transfected with pGC-PDK2^N255A^ or pGC^WT^ and acetate or PBS ctl treatement. N = 3 biological replicates. **h, i.** Representative western blot (h) and quantification (i) of H3K27ac levels in cNFs transfected with siCtl/siPDK2 and c646 or DMSO ctl treatment. N ≥ 3 biological replicates. β-tubulin was used as a loading control. **j, k.** Representative western blot (j) and quantification (k) of H3K27ac levels in pNF2 transfected with siCtl/siPDK2 and c646 or DMSO, as control, treatment. N ≥ 3 biological replicates. β-tubulin was used as a loading control. **l, m.** mRNA expression of *COL1A1*, *COL6A1* and *PYCR1* in cNFs (l) and pNF2 (m) transfected with siCtl/siPDK2 and treated with c646 or DMSO, as control. N ≥ 3 biological replicates. **n.** Quantification of COL6A1 in decellularised ECM derived from pNF transfected with siCtl/ and treated with c646 or DMSO, as control. N = 3 biological replicates. **o, p.** Representative western blot (o) and quantification (p) of COL6A1 in decellularised ECM derived from cNFs transfected with siCtl, siPDK2 or siPDK2 + siPYCR1. N = 3 biological replicates. **q, r.** Representative western blot (q) and quantification (r) of COL6A1 in ECM derived from pNF2 transfected with siCtl, siPDK2 or siPDK2 + siPYCR1. N = 3 biological replicates. Error bars indicate mean ± SEM. *p ≤0.05, **p ≤0.01, ***p ≤0.001. See Extended Data Figure 9 for red Ponceau staining of the blots used for COL6A1 staining in the ECM, which was used to normalise to total protein content in each lane.

Finally, we assessed the requirements of PYCR1 to support increased collagen production following increased PDC activity. For this, we silenced PDK2 in NFs, alone or together with PYCR1 (**Extended data Figure 7m, n**), and monitored collagen production. Strikingly, reducing PYCR1 levels in NFs silenced for PDK2 was sufficient to inhibit collagen production induced by PDC activation (**Figure 8p-r**). Hence, PDC activity is a key regulator of collagen production in CAFs and targeting PYCR1 effectively opposes this function.

## Discussion

Our work provides the first evidence for a link between proline biosynthesis and tumour ECM production *in vivo*, as we show that, in mice subcutaneously transplanted with MCF10DCIs.com xenografts, tumours use circulating glutamine to generate proline for integration into tumour collagen. Using different models of human mammary NF and CAF that produce abundant collagen, we uncovered proline synthesis via PYCR1 as major regulator of enhanced collagen production. Collagen has been shown to have both tumour promoting and tumour protecting functions ^34–42^. Targeting PYCR1 in CAFs in co-transplantation models of breast cancer reduced tumour collagen and was sufficient to reduce tumour growth and metastasis. Our work adds to the increasing body of evidence that cellular metabolism supports pro-tumorigenic traits of CAFs ^51–58, 96^ and identified PYCR1 as a candidate target to reduce tumour collagen to oppose breast cancer progression. It remains to be seen whether this is also the case when heterogeneous populations of CAF populate the tumour stroma and in other tumour types, such as PDAC, in which CAFs and collagen have been shown to have tumour-restraining functions ^6, 41, 45, 46^.

Amino acid metabolism in CAFs influences tumour incidence and metastatic spread because cancer cells can develop dependencies on amino acids produced and secreted by CAFs to fuel their growth ^51–57^. For example, tumour collagen-derived proline promotes PDAC cell survival under nutrient limited conditions ^56^. Here we establish that proline is also important to support intrinsic CAF functions. In fact, we provide evidence that enhanced proline synthesis through PYCR1 is a metabolic requirement for biosynthesis of abundant collagen-rich ECM (**Extended Data Figure 8**). PYCR1 activity can be regulated by post-translational modifications ^97^, and while we show that increased proline synthesis is achieved via increasing PYCR1 levels, we cannot exclude that also its activity is higher in CAFs. Our findings are consistent with the knowledge that collagens are proteins with an exceptional high proline content ^61^, and that fibroblasts in culture can generate proline from glutamine through PYCR1 ^63, 64, 98–100^. Interestingly, Schwörer and colleagues ^100^ recently showed that targeting PYCR1 in fibroblasts did not affect collagen production induced upon TGFβ-stimulation. In contrast, ALDH18A1 ablation reduced collagen synthesis, because its activity requires ATP and NADH, thus enabling the quenching of the high mitochondrial redox potential generated by TGFβ-mediated stimulation of glucose and glutamine metabolism ^100^. This suggests that under acute TGFβ stimulation, fibroblasts increase collagen production and secretion to protect themselves from oxidative damage. Whether the same mechanism is active after fibroblasts have been fully activated into myofibroblasts has not been addressed by the authors. In our NF and CAF models, we could not find consistent differences in glucose and glutamine usage and targeting PYCR1 in CAFs resulted in a reduction of collagen in the stroma of tumour xenografts and in 2D and 3D co-cultures with cancer cells. This suggests that increased collagen production in CAFs may not be a consequence of the redox imbalance described in acutely TGFβ-activated fibroblasts. In our work, we did not investigate whether PYCR1 controlled the redox status of CAFs. However, as silencing PYCR1 affected ribosomal stalling at proline codons of several collagens and we could rescue collagen translation and ECM production by providing exogenous proline, we propose that a major function of the proline synthesis pathway in CAFs is to provide proline residues for ECM production. Recently Guo et al. uncovered that PYCR1 levels in cancer cells may also play a role in ECM production in tumours. The authors showed that deletion of kindlin 2 or PINCH1 in cancer cells of a lung adenocarcinoma model led to decreased levels of PYCR1 and proline, and that these correlated with significant reductions of tumour growth and collagen in the ECM ^101, 102^.

Importantly, we found that acetyl-CoA is another major metabolic regulator of collagen production in CAFs. Acetyl-CoA is known to be an epigenetic modifier in cancer and immune cells through influencing histone acetylation, and EP300 is a key mediator ^103, 104^. Our work shows that increasing acetyl-CoA levels in fibroblasts induced H3K27 hyperacetylation and a transcriptional rewiring with elevated expression of pro-tumorigenic collagens, *COL1A1*, *COL6A1* ^38–40^, and *PYCR1*, and that EP300 is required for this process. Fibroblasts and CAFs can be epigenetically reprogrammed by DNA and histone methylation ^54, 57, 86^. Recently Eckert et al. have shown that targeting the methyl transferase nicotinamide N-methyltransferase (NNMT) in CAFs influences histone methylation and reduces tumour growth and metastatic potential ^57^. Our work reveals acetyl-CoA as another major epigenetic regulator in CAFs. Whether increased acetyl-coA in CAFs induces hyperacetylation of histones other than H3K27, and whether it regulates H3K27 acetylation solely by providing the acetyl group substrate, or it also acts as a signalling molecule, for example by regulating EP300 activity and selectivity ^105^, is still an open question. Similarly to fibrosis, we also found that *COL1A1* and *COL6A1* genes are under the control of EP300 in CAFs^106^ and further discovered that *PYCR1* is co-regulated and necessary to support the biosynthetic requirement for increased collagen synthesis. Previous work has shown that *PYCR1* expression is regulated by the amino-acid starvation response (AAR/*Atf4*) ^107^ and induced by shortage of proline precursors ^74^. We speculate that in CAFs EP300 directly controls collagen expression, while *PYCR1* expression increases in response to subsequent reduction in proline availability due to enhanced collagen synthesis. However, further studies are needed to tackle this yet unanswered question.

Another novel finding of our study is that PDK2 is the most highly expressed of the four PDK isoenzymes in mammary NFs and major regulator of PDC activity, acetyl-CoA production and gene expression rewiring in mammary CAFs and NFs. Since PDC-derived mitochondrial acetyl-CoA requires ACLY to contribute to the pool of nucleo-cytosolic acetyl-CoA, this implies ACLY as an additional epigenetic regulator in CAFs. This is supported by our observation that pharmacological inhibition of ACLY reduced acetyl-CoA levels and H3K27 acetylation, and that this correlated with reduced proline synthesis and production of collagen. Due to the challenge of performing metabolic tracing experiments *in vivo*, particularly in stromal cells that constitute only a small portion of the tumour, we have not validated *in vivo* whether, similarly to the *in vitro* setting, PDC-activity is a major source of acetyl-CoA in CAFs. However, our analysis of gene expression in stroma dissected from normal and tumour tissue showed that *PDK2* expression is also reduced in primary breast cancer patient samples. Moreover, previous works reported decreased levels of PDK proteins in the stroma of patient-derived non-small cell lung cancer samples compared to normal lungs ^108, 109^, supporting the conclusion that PDC activity is enhanced in the stroma of malignant tumours. Future experiments are needed to confirm this hypothesis and to assess whether other metabolites contribute to the modulation of acetyl-CoA levels in CAF *in vivo*, such as acetate through ASSC2 or glutamine through reductive carboxylation. This is particularly relevant under low nutrient availability and hypoxia, since these stress conditions increase ACSS2 expression and acetate usage to make acetyl-CoA in cancer cells ^110–112^, and in tumours in which cancer cells can synthesise and secrete acetate ^113^, which, in turn, may be used as metabolic intermediate by neighbouring cells ^114^. Interestingly, while supraphysiological levels of acetate were not able to significantly alter H3K27 acetylation or ECM production in CAF *in vitro*, they were sufficient to do so in normal fibroblasts. High levels of circulating acetate occur in individuals with chronic ethanol consumption, in which plasma acetate concentrations can reach above 0.8 mM^115^. Further studies are warranted to assess whether this may influence ECM production that underlines fibrotic (and ultimately cirrhotic) liver disease associated with alcohol consumption.

Throughout our work we have used models of mammary CAFs with enhanced ECM production compared with their NF counterpart and found *PYCR1* and *COL1A1* both expressed at higher levels in LCMD IDC stroma and in CAFs with a myofibroblast phenotype in scRNAseq analyses of human TNBCs. Together, these results suggest that proline synthesis via PYCR1 for collagen production is an acquired trait of fibroblasts to support abundant ECM production. Whether enhanced de novo proline synthesis via PYCR1 is a common feature of activated fibroblasts and required for collagen production in other tumour types and its role in tumour progression has to be determined.

In conclusion, our work uncovered that proline metabolism provides a critical link between the epigenetic regulation of collagen gene expression and protein production in CAFs, thus offering a new paradigm, whereby CAF metabolism emerges as major vulnerability of pro-tumorigenic collagen production. This is an important finding because *PYCR1* is among the top 20 metabolic genes overexpressed across cancer types ^116^, and proline synthesis has been proposed as a tumour-specific vulnerability ^117–119 52, 85, 99, 120^. Here we show that *PYCR1* and collagen upregulation co-occur in many types of tumours, hence we anticipate that further work exploring PYCR1 as a therapeutic target in tumours, to attack both cancer and stromal cells, may result in development of novel strategies to treat cancer. Moreover, collagen is the major contributor to the formation of desmoplastic tumour stroma. Thus, our study implies that targeting PYCR1 may also offer opportunities to tackle tumour-associated fibrosis to improve effective drug delivery and immune cell recruitment.

## Methods

### Cell culture

Patient-derived mammary cancer-associated fibroblasts and normal fibroblasts (pCAFs and pNFs) were isolated in house from breast cancer patient samples obtained through NHS Greater Glasgow and Clyde Biorepository. All participants gave specific consent to use their tissue samples for research. The cancer cell-derived, immortalised, GFP positive human mammary CAFs and NFs (cCAFs and cNFs) were kindly provided by Professor Akira Orimo (Juntendo University, Tokyo). The expression of αSMA between paired NFs and CAFs was routinely monitored to ensure that during our experiments the NFs are not activated into myofibroblasts. The fibroblasts and HEK293T cells were cultured in Dulbecco’s modified Eagle’s medium (DMEM) supplemented with 10% foetal bovine serum, 2 mM glutamine and 1% penicillin/streptomycin. For physiological DMEM (Physiol DMEM) experiments, fibroblasts were cultured in DMEM containing 5 mM glucose and supplemented with 0.65 mM glutamine, 100 µM pyruvate, 100 µM acetate, 10% FBS and 1% penicillin/streptomycin. MCF10DCIS.com cells were cultured in F12 medium supplemented with 5% horse serum, 2 mM glutamine, 1% penicillin/streptomycin and 0.1% fungizone. Wood primary breast cancer cells were purchased from AMS Biotechnology Europe Ltd (AMSBIO) and cultured in Renaissance essential tumour medium (RETM, AMSBIO) supplemented with 5% fetal bovine serum and 1% penicillin/streptomycin. For 2D and 3D co-cultures, CAFs and cancer cells were mixed in a 1:1 ratio and cultured in a 1:1 mixture of DMEM and RETM. For SILAC proteomics experiments, iCAFs and iNFs were cultured in SILAC DMEM supplemented with 2% FBS, 8% 10 kDa dialysed FBS (PAA), 2 mM glutamine and 1% penicillin/streptomycin. SILAC DMEM used for the ‘light’ labelled cells contained 84 mg/l L-arginine and 146 mg/l L-lysine (Sigma), whereas the medium for the ‘heavy’ labelled cells contained 84 mg/l ^15^N L-arginine and 175 mg/l ^13^C_6_^15^N_2_ L-lysine (Cambridge Isotope Laboratories). Cells were regularly tested for mycoplasma and MCF10DCIS.com cells authenticated. The following inhibitors were used to treat cells in culture: c646 (Sigma), A485 (Tocris Bioscience), BMS303141 (Sigma) and CPI-613 (Sigma). The PYCR1i was made as previously described ^71^. Unless otherwise stated, cells were treated with compounds at the following concentrations: 25 µM c646, 3 µM A-485, 50 µM BMS30341, 100 µM CPI-613, 20 µM PYCR1i, 1 mM acetate, 500 µM proline.

### pCAF and pNF isolation and immortalisation

pCAFs and pNFs were isolated in house from patient samples using previously described methods ^22, 25, 59^. From each patient, pCAFs were isolated from breast tumour tissue and pNFs from normal, tumour adjacent tissue. Patient samples were obtained complying with ethical regulations through the National Health Service (NHS) Greater Glasgow and Clyde Biorepository. All participants gave specific consent to use their tissue samples for research. pCAF/NF2 were from a ER+, PR+, HER2-breast cancer patient and pCAF/NF3 were from a triple negative breast cancer patient. The pCAFs and pNFs were immortalised using a human telomerase reverse transcriptase (hTERT)-expressing plasmid (pIRES2-hygro), kindly provided by Dr. Fernando Calvo (IBBTEC, Santander). Lentivirus containing the hTERT plasmid was generated in HEK293T cells. Two rounds of viral transduction in fibroblasts were performed on consecutive days. Cells were selected using 50 μg/ml hygromycin.

### Western blotting analysis

Cells were cultured for 48h, following transfection and/or treatment with inhibitors and rescue compounds, then lysed in SDS buffer (2% SDS, 100 mM TrisHCl pH 7.4), incubated at 95 °C for 5 min, sonicated using a metal tip (Soniprep 150, MSE) and centrifuged at 16000 x g for 10 min. Protein concentration was determined using Optiblot Bradford reagent (Abcam). 20-25 μg of proteins were separated using 4-12% gradient NuPAGE Novex Bis-Tris gel (Life technologies). Protein transfer was performed on methanol-activated PVDF or Nitrocellulose membrane. Membrane was blocked for 1 hr in 3% BSA (Sigma) in TBST at RT and incubated with primary antibodies overnight at 4°C. The following primary antibodies were used: PDHA1 E1-alpha subunit, phospho S293 (1:2000, abcam ab177461), PDHA1 E1-alpha subunit (1:1000, abcam ab110334), PYCR1 (1:1000, Proteintech 22150-1-AP), Histone H3, acetyl K27 (1:1000, abcam ab4729), Collagen VI (1:1000, abcam ab182744). Vinculin (1:2000, Sigma V9131), GAPDH (1:1000, Santa Cruz, sc48167), β-tubulin (1:1000, abcam ab179513) were used as loading controls for the experimental antibodies. The membrane was incubated with HRP-conjugated secondary antibodies (1:5000, NEB) for 45 mins at RT. Western blot images were acquired using a myECL Imager (Thermo Scientific).

### PDH activity assay

PDH activity was measured using the Pyruvate dehydrogenase Enzyme Activity Microplate Assay Kit (abcam ab109902) according to the manufacturer’s protocol.

### EDU proliferation assay

Cells were seeded on 13 mm glass coverslips. Following 48 h of drug treatment, cells were incubated in 1 µM EDU for 2 h and fixed in 4% PFA. EDU was fluorescently labelled using the Click-iT™ EdU Cell Proliferation Kit (Life Technologies) according to the manufacturers’ protocol, and nuclei were counterstained with DAPI. Images were acquired using a Zeiss 710 confocal microscope and ImageJ was used to count the number of total nuclei and EDU positive nuclei.

### Decellularised ECM preparation

Cells were seeded at 100% confluence on 0.2% gelatine, which was crosslinked using 1% glutaraldehyde, then were cultured either for 7 days with inhibitor treatment or 3 days if they had undergone transfection with siRNA or plasmids to ensure the construct was expressed for the duration of the experiment. The ECM was decellularised with extraction buffer (20mM NH_4_OH, 0.5% Triton X-100 in PBS) until no intact cells were visible but the ECM remained on the dish. ECM was washed in PBS with Ca^2+^ and Mg^2+^, collected and lysed in SDS buffer (4% SDS, 0.1 M DTT, Tris-HCl pH 7.4).

### Cell transfection and infection

For transient expression or siRNA knockdown, 2×10^6^ fibroblasts were harvested and used in each transfection with a Nucleofector device (Lonza) according to the manufacturer’s protocol using the program T-20 and the Amaxa kit R (Lonza). Cells were transfected with 1-3 nM non-targeting siRNA as a control (D-001810-10-05, GE Healthcare Dharmacon) or with siRNAs targeting *PDK2* and *PYCR1* (Dharmacon, pool of 4), or with 5µg pGCA-PDK2^N255A^ or pGCA-PDK2^WT^ (kindly provided by Prof. Angus McQuibban, University of Toronto ^47^). Cells were used for experiments 48-72h after transfection.

For stable knock down of PYCR1, shPYCR1 (shPCYR1 #1: CCGGTGAGAAGAA GCTGTCAGCGTTCTCGAGAACGCTGACAGCTTCTTCTCATTTTTG, shPYCR1 #2: CCGGCACAGTTTCTGCTCTCAGGAACTCGAGTTCCTGAGAGCAGAAACTGTGTTTTTG) and shCTL (Sigma, Mission shRNA) lentivirus was generated in HEK293 cells. Two rounds of viral transduction in pCAF2 were performed on consecutive days. Cells were selected using 2 µg/ml puromycin.

### Reverse transcriptase polymerase chain reaction (RT-qPCR)

Primers for RT-qPCR were designed using Primer Blast (NCBI database): *PDK2*: CGGGGACCACAACCAAAGTC (forward) GCTGGATCCGAAGTCCAGAAA (reverse), *PYCR1*: CCCCGCCTACGCATTCACA (forward) GCGCGTTGGAAGTCCCATCT (reverse), *COL1A1*: TGAAGGGACACAGAGGTTTCAG (forward) GTAGCACCATCATTTCCACGA (reverse), *COL6A1*: AGCAAGTGTGCTGCTCCTTC (forward) CTTCCAGGATCTCCGGCTTC (reverse). Akaluc construct. Data was normalised to expression of the following housekeeping genes: *TBP2*: AGTGACCCAGCATCACTGTTT (forward) TAAGGTGGCAGGCTGTTGTT (reverse), *18S*: AGGAATTGACGGAAGGGCAC (forward) GGACATCTAAGGGCATCACA (reverse) or *ACTB*: GGCATGGGTCAGAAGGATT (forward) ACATGATCTGGGTCATCTTCTC (reverse). For gDNA extraction from whole blood, red blood cells were lysed in a hypotonic solution of 0,2% NaCl for 30 s and brought in isotonic condition with 1.6% NaCl and 0.1% glucose. The remaining leukocytes and circulating tumour cells were washed once with PBS and gDNA was extracted from the pellet using the QIAmp DNA mini kit (Qiagen). Total RNA was isolated from cells after 48h in culture, following transfection and/or with inhibitor and rescue compound treatment. RNA was isolated with the RNEasy mini kit (Qiagen) according to the manufacturer’s instructions. Complementary DNA (cDNA) was synthesised from 1 µg RNA using an iScript kit (BioRad). DNA was diluted to 10 ng/µland 2 µl was used in each RT-qPCR reaction with 10 µl iTAQ Universal SYBR green supermix (BioRad) and 400 nM primers. Reactions were performed using a Quant Studio 3 PCR machine (Thermo Scientific).

### MS-proteomic analysis

For the total proteome, cells were lysed in SDS buffer (2% SDS, 100 mM TrisHCl pH 7.4). For the proteome with c646 treatment, proteins were precipitated with acetone and redissolved in urea buffer (6M urea, 2M thiourea, 10 mM TCEP, 40 mM CAA, 75 mM NaCl, 50 mM Tris-HCl). The proteins were then trypsin digested. For SILAC experiments, equal quantities of heavy and light samples were mixed. For the total proteome that was used to normalise peptide acetylation levels, lysates were either in-gel digested with trypsin or, after trypsin digestion, peptides were fractionated using high pH reverse phase fractionation. The proteins were desalted by C18 StageTip ^121^ prior to MS analysis.

For phosphorylated peptide enrichment, trypsin-digested peptides were acidified to pH 2.6 and acetonitrile (ACN) was added to a final concentration of 30%. The peptides were fractionated using an Akta system into 6 equal fractions, using an increasing concentration of KCl in 5 mM KH_2_PO_4_ to a final concentration of 350 mM KCl. Each fraction was then enriched for phosphorylated peptides by incubation with TiO_2_ beads (GL Sciences) in the presence of 2,5-dihydroxybenzoic acid ^122^. Phosphorylated peptides were eluted with 15% ammonium hydroxide and 40% acetonitrile (ACN), and desalted by C18 StageTip.

For acetylated peptide enrichment, the deacetylase inhibitors nicotinamide (10 mM) and trichostatin A (1 µM) were added to the lysis buffer. Different protocols were used to prepare acetylated peptides. i) Cells were lysed in RIPA buffer (50 mM TrisHCl pH 7.5, 150 mM NaCl, 1 mM EDTA, 1% NP-40, 0.1% sodium deoxycholate). Proteins were precipitated with acetone, redissolved in urea buffer and quantified by Bradford assay. Equal quantities of heavy and light labelled proteins were combined and trypsin digested. Peptides were desalted by C18 SepPak filtration and resuspended in immunoprecipitation (IAP) buffer (50 mM MOPS; pH 7.2, 10 mM Na-phosphate, 50 mM NaCl). Acetylated peptides were enriched using anti-acetyllysine antibody (Acetyl Lysine Antibody, Agarose, ImmuneChem). Up to three consecutive incubations were performed to maximise peptide recovery. Acetylated peptides were eluted with acidified water (0.1% TFA in water). ii) Lysis and protein digestion was performed as in i), but the enrichment for acetylated peptides was performed with PTMScan Acetyl-Lysine Motif Kit (Cell Signalling Technology #13416) according to the manufacturers’ protocol. iii) Subcellular fractionation of the cells was performed (Cell Fractionation Kit, Standard, Abcam, ab109719) according to the manufacturers’ protocol. Proteins recovered in the three fractions, nuclear, cytosolic and mitochondrial, were digested with trypsin and acetylated peptides enriched using anti-acetyllysine antibody (Acetyl Lysine Antibody, Agarose, ImmuneChem).

For ECM analysis, cell-derived ECM was prepared as above. To analyse glutamine-derived proline incorporation into collagen, the cells were cultured in media containing 2 mM ^13^C_5_-glutamine for 72 h prior to ECM collection. Each sample was separated on 4–12% gradient NuPAGE Novex Bis-Tris gel (Life Technologies). The gel was sliced into 3 fractions, and each fraction was in-gel digested with trypsin.

For proteomic analysis of tumours, 100 µg tissue was homogenised in 4% SDS, 0.1 M DTT buffer. The resulting lysate was precipitated with acetone, re-dissolved in urea buffer, and trypsin digested. Peptides were resuspended in 1% TFA, 0.2% acetic acid or formic acid buffer and injected on an EASY-nLC (Thermo Fisher Scientific) coupled online to a mass spectrometer. Peptides were separated on a 20-cm fused silica emitter (New Objective) packed in-house with reverse-phase Reprosil Pur Basic 1.9 µm (Dr. Maisch GmbH). Peptides were eluted with a flow of 300 nl/min from 5% to 30% of buffer B (80% ACN, 0.1% formic acid) in a 60-min linear gradient. Eluted peptides were injected into an Orbitrap Elite, Q-Exactive HF or Orbitrap Fusion Lumos (Thermo Fisher Scientific) via electrospray ionisation. MS data were acquired using XCalibur software (Thermo Fisher Scientific).

### MS-proteomic data analysis

The MS .raw files were processed with MaxQuant software ^123^ and searched with the Andromeda search engine with the following settings: minimal peptide length 7 amino acids, fixed modification Carbamidomethyl (C) and variable modifications Acetyl (Protein N-term) and Oxidation (M). For the acetylome and phosphoproteome, Acetyl (K) and Phospho (STY) were added as variable modifications respectively. For the tracing experiments with ^13^C_5_-glutamine, proline oxidation was added as a variable modification. For SILAC experiments, multiplicity was set to 2, where the light labels were Arg0 and Lys0 and the heavy labels were Arg10 and Lys8. For the tracing experiments with ^13^C_5_-glutamine, ^13^C_5_-proline and ^12^C_5_-proline were added as heavy and light labels respectively. For label free quantification (LFQ) experiments, the LFQ setting was enabled. The false discovery rates (FDRs) at the protein and peptide level were set to 1%. Specificity for trypsin cleavage was required and maximum 2 missed cleavages were allowed.

Perseus ^124^ (version 1.5.0.36 for the phosphoproteome, 1.5.5.1 for the acetylome and corresponding proteome, 1.6.2.2 for total proteome) was used for downstream analysis. The data were filtered to remove potential contaminants, reverse peptides which match a decoy database, and proteins only identified by site. To ensure unambiguous identification, only proteins identified with at least one unique peptide were considered. For SILAC experiments, the SILAC ratio was used for the analysis. Ratios from the ‘Reverse’ experiment were inverted. Then, SILAC ratios were transformed by log2 and intensities by log10. For the acetylome, peptide acetylation levels were normalised by total protein abundance (**Supplementary Data S2**). For ^13^C_5_-glutamine tracing experiment). For ^13^C_5_-glutamine tracing experiments, heavy (^13^C-proline) and light (^12^C-proline) labelled peptides were filtered as described above and transformed by log_2_.

### Estimation of kinase activities

KinAct ^125^, is a computational method used to predict kinase-activity scores from MS-based data. It infers an activity score for each protein kinase based on the regulation levels of phosphorylation events catalysed by this specific kinase. The method relies on prior knowledge of kinase/phosphatase-to-substrate (k/p-s) relations and the KSEA (Kinase-Substrate Enrichment Analysis) method ^126^ for the kinase activity estimation.

Currently, there are multiple freely available databases that collect k/p-s interactions which are experimentally verified or manually curated. Some of them are integrated into OmniPath ^127^, a comprehensive collection of signalling resources. There are also resources in which the k/p-s relations are inferred based on consensus kinase recognition motifs and other information, such as NetworKIN ^128^). The KSEA method integrates the information from such databases with the log_2_ transformed fold changes from mass-spectrometry data to compute enrichment scores for each kinase together with a significance value for each score.

In the KSEA method, the score is equal to the mean of the fold changes of each phospho-measurement of the substrate set *mS* of a specific kinase. The significance of the score, on the other hand, is calculated from a *z*-statistic as: 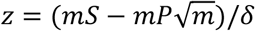, where *mP* is the mean log_2_ fold-change of the complete data-set, *m* is the size of the substrate set *mS* (how many substrates for a kinase) and *δ* is the standard deviation of the log_2_ fold-change values for the whole dataset. Significance is then estimated from the *z*-statistic and the estimated *p*-values for each kinase are then adjusted via the Benjamini-Hochberg correction method.

KinAct was applied to the phosphoproteomic SILAC-labelled NF (normal fibroblast) and CAF (cancer-associated fibroblast) data which was performed in two independent experiments. The log_2_ ratios of the two experiments were averaged and inputinto the KinAct pipeline. KinAct analysis was performed over the OmniPath and NetwroKIN prior knowledge. For the OmniPath prior knowledge, kinase-substrate relations coming from PhosphoSitePlus ^129^ and SIGNOR ^130^ were used. Results from both resources yielded similar results where PDK2 was observed to be significantly downregulated in both cases. Codes of the analysis together with usage documentation are made available in https://github.com/saezlab/iCAF_KinAct.

### Metabolites extraction and LC-MS analysis

For tracing experiments, cells were labelled for 24h with ^13^C_6_-glucose, ^13^C_5_-glutamine, ^13^C_3_-pyruvate (100 µM) or ^13^C_2_-acetate (100 µM). With the exception of acetate, which was supplemented to the media, the ^13^C-labelled metabolite replaced the concentration of the metabolite in DMEM. Cells were optionally treated with inhibitors/rescue compounds for the duration of the experiment. Cells were washed with ice-cold PBS and intracellular metabolites were extracted with extraction buffer (aqueous solution of 50% methanol and 30% acetonitrile). Blood samples were diluted 1:50 in extraction buffer and incubated with shaking at 4°C. Tumour samples were homogenised at 4°C in extraction buffer at a concentration of 20 mg/ml (tissue/extraction buffer). Extracts were centrifuged at 16,000 x g for 5mins, and the supernatant was analysed using a Q-Exactive Orbitrap mass spectrometer (Thermo Scientific) in combination with a Thermo Ultimate 3000 HPLC system. 5 μl of cell extract was injected and the metabolites were separated over a 15 min mobile phase gradient from an initial ACN content of 80% ACN with 20% ammonium bicarbonate (pH 9.2) decreasing to 20% ACN with a flow rate of 200 μL/min. The metabolites were detected over a period of 25 min using the Q-Exactive mass spectrometer across a mass range of 75-1000 m/z and at a resolution of 35,000 (at 200 m/z). To detect acetyl-coA, a single ion monitoring (SIM) method was employed. The Q-Exactive mass spectrometer was used to monitor the three masses for acetyl-coA labelled +0, +1 or +2 (810.1331, 811.1364 and 812.13976 m/z) with an isolation window of 0.7 m/z for each isotope. Peak identification and area quantification were carried out using TraceFinder software by comparison of the retention time and exact ion mass to that of authentic standards.

### Ribosome profiling and diricore analysis

30×10^6^ cells were treated with cycloheximide (100 μg/ml) for 5 minutes and lysed in buffer A (20 mM Tris-HCl, pH 7.8, 100 mM KCl, 10 mM MgCl_2_, 1% Triton X-100, 2 mM DTT, 100 μg/ml cycloheximide, 1X complete protease inhibitor). Lysates were treated with 2 U/μl of RNase I (Ambion) for 45 min at room temperature. Lysates were fractionated on a linear sucrose gradient and the fractions enriched in monosomes were pooled. Ribosome protected fragments (RPFs) were purified using Trizol reagent (Invitrogen). Library preparation and differential ribosome codon reading (diricore) analysis were performed according to the method previously described ^74^.

### Collagen quantification in mono culture and 2D co-cultures

Cells were seeded at 100% confluence on glass coverslips, either as a CAF monoculture or as a 1:1 coculture of CAFs and Wood primary breast cancer cells. The cells were cultured for 96h in the presence of inhibitors or rescue compounds to allow accumulation of matrix. Cells were incubated with 1 µM of the native collagen binding protein CNA35 labelled with fluorescent dye mCherry (CNA35-mCherry) ^73^ for 1 hr to label collagen, then fixed in 4% PFA and counterstained with DAPI. Images were taken on a Zeiss 710 confocal microscope. Regions of CAFs were defined and collagen staining was quantified using ImageJ software. Cell number for normalisation of collagen quantification was calculated by counting the number of DAPI positive nuclei using ImageJ software.

### Microfluidic device design and preparation

Microfluidic devices were fabricated using previously established soft lithography methods and used to culture spheroids ^75^. Multilayer devices were composed of arrays of microfluidic channels, each of which was connected by two open wells. Within each channel, an array of microwells of dimension 150×150×150 µm^3^ were situated below the channel level ^75^. In short, polydimethylsiloxane (PDMS) prepolymer (Sylgard 184, Dow Corning) and curing agent were combined in a 1:10 ratio and poured onto patterned silicon wafers. Wafers were placed inside a desiccator for degassing, prior to incubation at 85°C for a minimum duration of 3 hours. Once cured, the PDMS was removed from the wafers and open wells created using a 4mm surgical biopsy punch (Miltex). Devices were cleaned and exposed to an oxygen plasma (Pico plasma cleaner, Diener electronic) to permanently bond the upper and lower PDMS layers together. Devices were incubated with a solution of 1% Synperonic F108 solution (Sigma Aldrich) to achieve ultra-low adhesion conditions.

### 3D co-culture in microfluidic devices

Cells were seeded at a 1:1 ratio of Wood primary breast cancer cells:CAFs into devices at a concentration of 7×10^6^ cells/ml to form spheroids, with each microfluidic channel containing at least 32 spheroids of similar dimension (∼80 µm diameter) for analysis. A cell suspension was injected in the open wells, which flowed into the microfluidic channels until they remained trapped into the microwells, as previously described ^75^. Spheroids were formed within 24-48 hours. Cells that were not trapped into the microwells were removed from the device. Cells were cultured in a 1:1 mix of the supplier recommended complete culture media for the two cell types: RETM for the primary breast cancer cells and DMEM for the CAF. Media with and without enzyme inhibitors and rescue agents was exchanged every 48 hours.

### Enzyme inhibitor treatment in microfluidic devices

Spheroids were treated with BMS303141 (Concentrations: 25 µM, 50 µM and 100 µM), c646 (Concentrations: 25 µM, 50 µM and 100 µM), PYCR1i (Concentrations: 50 µM, 100 µM and 150 µM) or CPI-613 (Concentrations: 50 µM, 100 µM and 200 µM). Inhibitor action on cells was mitigated with administration of either 1 mM acetate or 500 µM proline, used as rescue agents and applied together with the enzyme inhibitor treatment. Inhibitor treatment was administered every second day for one week beginning 24 hours after cell seeding. Control experiments were performed for each experimental setup. All experiments were performed in triplicates, with each experiments performed on at least 32 spheroids.

### Collagen quantification in 3D co-cultures

For visualisation of total collagen, 1 µM CNA35-mCherry was incubated with the cells for a 2-hour period. After this, cells were washed twice with PBS to ensure removal of any residual staining solution. PBS was then added again prior to imaging the devices.

An inverted microscope (Observer A1, Zeiss) connected to an Orca Flash 4.0 camera (Hamamatsu) was used to acquire bright field images of spheroids every 24–48 hours. Epifluorescence microscopy was performed immediately after cell staining and image analysis carried out using ZEN Blue and Fiji. czi files from ZEN Blue were split into separate channels and converted into TIFF files in Fiji. Images from each channel were normalized to the same threshold range. Due to the distinct dissociation of the two cell types after 7 days of co-culture, it was possible to quantify the spheroid areas and perimeters of both CAFs (expressing green fluorescent protein) and the cancer cells (from bright field images). Collagen deposition was estimated from fluorescent images (mCherry) and was plotted as a ratio of collagen area vs CAF spheroid area. Signal intensity was measured in these regions correcting for background signal.

### MCF10DCIS.com-CAF xenograft

All mouse procedures were in accordance with ethical approval from University of Glasgow under the revised Animal (Scientific Procedures) Act 1986 and the EU Directive 2010/63/EU authorised through Home Office Approval (Project licence number 70/8645).

1.5×10^6^ pCAF2s expressing shCtl or shPYCR1 and 5×10^5^ MCF10DCIS.com in 200 µl 50% growth factor reduced phenol red free Matrigel (BD Biosciences) in PBS were injected subcutaneously into the flank of 8 week old female BALB/c nude mice (Charles River). Six mice per group were transplanted for each experiment. Mice were randomly allocated to the two groups. The mice were sacrificed 14 days after inoculation and the tumours were excised, weighed and fixed in 4% PFA. The tumours were sliced into 400 µm sections and Z-stacks of each section were captured. The collagen was imaged using second harmonic generation (SHG) microscopy in combination with confocal microscopy to detect GFP-expressing fibroblasts. Using an ImageJ macro, regions of human CAFs were defined for each slice of the Z-stack and the area of collagen surrounding the CAFs was quantified. For ^13^C_5_-glutamine tracing experiment, 1.5×10^6^ pCAF2 and 5×10^5^ MCF10DCIS.com in 200 µl 50% growth factor reduced phenol red free Matrigel (BD Biosciences) in PBS were injected subcutaneously into the flank of 8 week old female BALB/c nude mice (Charles River). 12 days after inoculation mice were injected intraperitoneally with 10 µl/g of 200 mM ^13^C_5_-glutamine and 6 with ^12^C_5_-glutamine. Mice were injected 6 times over 2 days, with the final injection 30 mins before sacrificing the mice. Blood was collected via cardiac puncture, and tumours and skin were frozen for metabolomic and proteomic analysis.

### Orthotopic 4T1 mammary tumor experiments

All mouse procedures were in accordance with ethical approval from the Institutional Animal Care and Research Advisory Committee of the K.U. Leuven. 4T1 cells were transduced with a lentiviral vector encoding Akaluc ^131^, a firefly luciferase analogue with higher sensitivity, puromycin resistance and miRFP670. Transduced cells were sorted according to miRFP670 expression and miRFP670+ cells hereby referred to as 4T1-Aka. 10^5^ 4T1-Aka cells and 10^6^ CAFs (shCtl or shPYCR1) were mixed in a volume of 50 μl of PBS and co-injected orthotopically to the right mammary fat pad of the second nipple of NMRI nu/nu female mice. To assess tumour hypoxia and tumour vessel perfusion, pimonidazole (60mg/kg, intraperitoneal) and fluorescein isothiocyanate (FITC)-conjugated lectin (Lycopersicon esculentum; Vector Laboratories; 0.05 mg intravenous) were injected into tumour-bearing mice 1 h or 10 minutes before tumour harvesting, respectively. At end stage, blood was collected by cardiac puncture and transferred to lithium heparin Microtainer Tubes (BD Biosciences). After perfusion of the mice with NaCl 0.9% through the right ventricle, tumours were collected and weighed. Samples were collected in PFA 2% for histological analysis or snap-frozen for metabolomic analysis.

### Lung colony formation assay

In brief, lungs were automatically digested in digestion buffer using gentleMACS octodissociator (Miltenyi Biotec). The digested lungs were strained with a 70μm strainer, centrifuged and plated in petri dishes with selection medium (RPMI supplemented with 10% FBS, 1% Q, 1% P/S and 60 μM of 6-thioguanine). At day 3 the selection medium was refreshed and at day 5 replaced with RPMI without 6-thioguanine. At day 6, colonies were fixed with PFA 4%, contrasted with 0.01% crystal violet in dH_2_O and counted.

### Histology analysis

The following antibodies were employed using an Agilent Autostainer Link 48; CD31 at 1/75 (Abcam, UK) and Hypoxyprobe Pimonidazole at 1/150 (Hypoxyprobe, USA). For Sirius red staining, rehydrated slides were stained for 2 hours in staining solution: equal volumes of 0.1% Direct red 80 (Sigma Aldrich) and 0.1% Fast green (Raymond A Lamb) combined in a 1:9 dilution with aqueous Picric acid solution (VWR).

Quantitative tissue analysis was performed on serial, formalin fixed paraffin embedded, mouse tumour sections using Halo software (v3.1.1076.363, Indica Labs). For Pecam1 and pimonidazole the region of interest (ROI) selected excluded any capsule around the tumour. For Sirius red, the quantification was performed both with a ROI including and excluding the tumour capsule The tumour capsule being defined as a collagen rich band of tissue of variable thickness which encapsulates the tumour in vivo. For each of the two studies, software parameters were set which most accurately defined the stain of interest and all sections within each study were analysed using the same settings. Sirius red, Pecam1 and Pimonidazole were given as a percentage area of the tumour with data normalised to the average of the control for each group.

### Gene expression analysis

The breast cancer (GSE90505) microarray datasets were downloaded from the Gene Expression Omnibus (GEO) using the R statistical environment, version 3.5.0, and the Bioconductor package GEOquery, version 2.40.0 ^132^. Differential gene probe expression analysis was done using the linear models and differential expression for microarray data (Limma) package version 3.29.8 ^133^.

Gene expression data from Ma et al. ^67^ have been downloaded from Oncomine ^134^. Probe g5902035_3p_a_at for *PYCR* and probe Hs.172928.0.A2_3p_a_at for *COL1A1* have been used for the analysis.

### TCGA data analysis

TCGA data from the Pan Cancer Atlas study available in cBioportal were used for the analysis, and were analysed with tools available in cBioportal ^68, 69^. For each tumour type, the quartiles for the *PYCR1* and *COL1A1* genes were calculated based on the mRNA expression Z-scores relative to diploid samples. Tumours defined as *PYCR1* and *COL1A1* high were those that expressed the two genes at levels within their respective 4^th^ (upper) quartile, while those define as *PYCR1* and *COL1A1* low were those that expressed the two genes at levels within the 1^st^ (lower) quartile. Disease specific survival curves for these two groups of samples were visualised in cBioportal, exported as PDF files and their format edited with the Illustrator software. Logrank Test p-values reported in the figure are those calculated in cBioportal and displayed on the plot with the survival curves.

### scRNA sequencing

Data were analysed as described in the original manuscript from Wu et al. ^11^. UMAPs were generated with Seurat version 4.0.2 and cell types annotated according to Wu et al.

### Calculation of amino acid content in ECM proteins

To calculate the amino acid composition of the human proteome, we retrieved all proteoforms from the Swiss-Prot section of UniProt database (release 2020_01). Lists of matrisome proteins were downloaded from the Matrisome Project (http://matrisomeproject.mit.edu/other-resources/human-matrisome/) ^60^ and used to annotate the proteins. Specifically, the “Core Matrisome” dataset was utilized which comprises glycoproteins, collagens and proteoglycans. We exploited the Biopython module ^135^ and developed a custom python script to calculate the amino acid frequencies in the entire proteome, the full matrisome as well as the individual core matrisome subcategories.

### Statistical analysis

GraphPad Prism version 7.0 was used for statistical analysis. For qPCRs, metabolomics data and in vivo data analysis data were tested for normality and for experiments with two conditions, a two-tailed unpaired t-test with Welch’s correction was used to determine the p-value, while for experiments with more than two conditions, a one way ANOVA test with Dunnett’s multiple comparison test was used. When data were not normally distributed, and for all other experiments, Mann-Whitney test (two samples) and Kruskal-Wallis with Dunn’s post hoc test (multiple comparison) were used instead. P ≤ 0.05 was considered significant. All graphs show the mean ± SEM of at least 3 biological replicates (independent experiments) unless otherwise stated. For MS-proteomic analysis, Perseus software was used for statistical analysis. A one-sample t-test for SILAC experiments or a 2-sample t-test for LFQ experiments was used to determine significantly regulated proteins and enable data visualisation as a volcano plot.

## Data availability

The .raw MS files and search/identification files obtained with MaxQuant have been deposited to the ProteomeXchange Consortium (http://proteomecentral.proteomexchange.org/cgi/GetDataset) via the PRIDE partner repository ^136^ with dataset identifier PXD018343 (Login details for reviewers are Username: and Pwd: rbvaJx0O) and PXD024746 (Login details for reviewers are Username: and Pwd: vtSVavh6). All unique materials used are readily available from the authors.

## Supplemental Data

**Supplemental Data S1. Amino acid count of the human proteome**

**Supplemental Data S2. Ranking of proteins identified in the proteome of SDS soluble cCAF ECM**

**Supplemental Data S3. Proteomic analysis of pCAF2-MCF10DCIS.com tumours and skin from ^13^C-glutamine infused mice**

**Supplemental Data S4. COL1A1 peptides carrying ^13^C-labelled proline in cCAF siCtl and siPYCR1**

**Supplemental Data S5. Acetylome and total proteome of mammary cCAF and cNF**

**Supplemental Data S6. Proteome cCAF treated with c646**

**Supplemental Data S7. Phosphoproteome of mammary cCAF and cNF**

**Supplemental Data S8. Prediction of activated and de-activated kinases in cCAF compared to cNF**

**Extended Data Figure 1.**
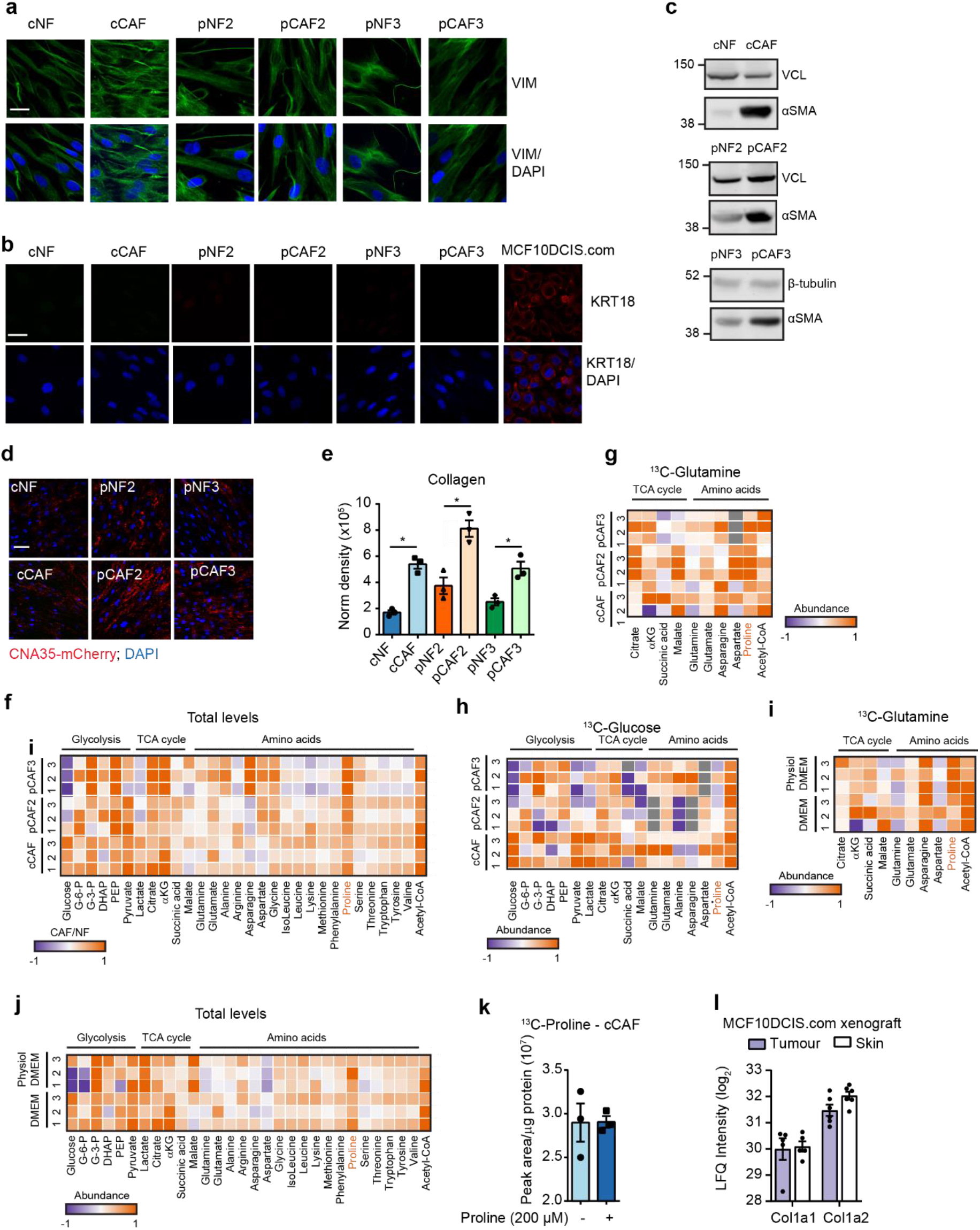
Characterisation of mammary CAF/NF pairs. **a.** Representative confocal microscopy images of vimentin staining in mammary CAF and NF pairs. Scale bar = 20 µm. **b.** Representative confocal microscopy images of keratin 18 (KRT18) staining in mammary CAF and NF pairs and MCF10DCIS.com breast cancer cells. Scale bar = 20 µm. **c.** Representative western blots showing α-SMA levels in mammary NFs and CAFs. **d, e.** Representative images (d) and quantification (e) of collagen produced by mammary NFs and CAFs. Scale bar = 50 µm. N = 3 biological replicates. **f.** Fold change in ^13^C-glutamine labelled glycolysis, TCA cycle and amino acid metabolites between paired CAFs and NFs. N = 3 biological replicates. **g.** Fold change in ^13^C-glucose labelled glycolysis, TCA cycle and amino acid metabolites between paired CAFs and NFs. N = 3 biological replicates.. **h.** Fold change in total levels of glycolysis, TCA cycle and amino acid metabolites between paired CAFs and NFs. N = 3 biological replicates. **f. i.** Fold change in ^13^C-Glutamine labelled metabolites between cCAFs cultured in standard DMEM with ^13^C-Glutamine or physiological DMEM (5 mM glucose, 0.65 mM glutamine, 100 µM pyruvate, 100 µM acetate) with ^13^C-Glutamine measured by MS-metabolomics. N = 3 biological replicates. **j.** Fold change in total levels of metabolites between cCAFs cultured in standard DMEM or physiological DMEM, measured by MS-metabolomics. N = 3 biological replicates. **k.** Total ^13^C-labelled proline in cCAFs cultured in physiological DMEM with ^13^C-Glutamine ± 200 µM proline, measured by MS-metabolomics. N = 3 biological replicates. **l.** Murine Col1a1 and Col1a2 protein levels in the tumour and skin tissues of mice carrying MCF10DCIS.com xenografts. Error bars indicate mean ± SEM. *p ≤0.05.

**Extended Data Figure 2.**
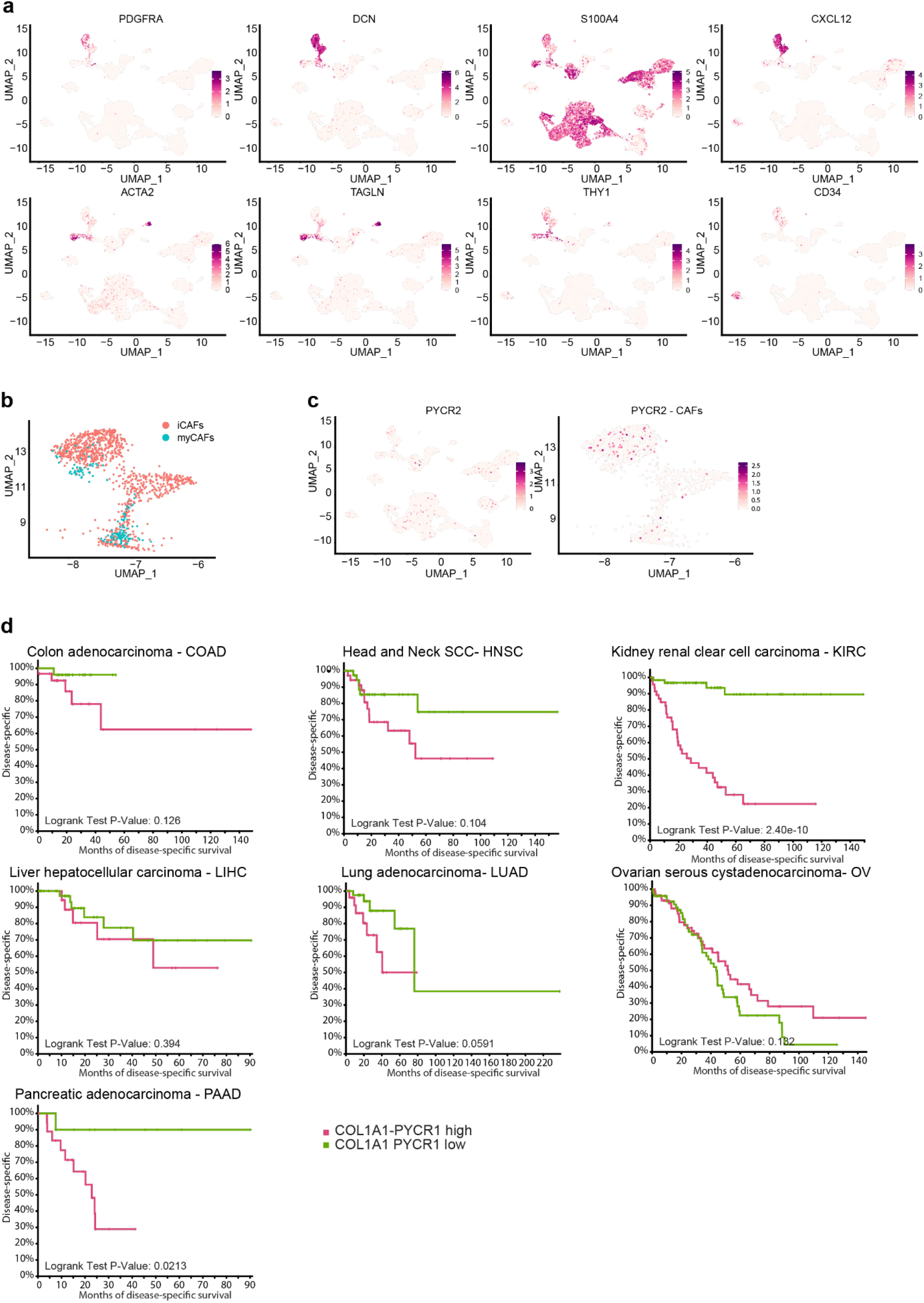
PYCR1 and COL1A1 expression in CAFs and cancer patients. **a.** UMAP visualisation of stromal, immune and cancer cells aligned using canonical correlation analysis in Seurat. Cell types defined as in Wu et al. **b.** UMAP visualisation of CAFs, with iCAF and myCAF as defined in Wu et al. **c.** UMAP visualisation of CAFs, as defined in Wu et al., aligned using canonical correlation analysis in Seurat. **d.** Kaplan-Meier plots comparing the disease specific survival of patients with tumours expressing high (top quartile) or low (bottom quartile) levels of both *COL1A1* and *PYCR1*. Data generated with cBioportal using TCGA Pan Cancer Atlas for the indicated tumour types.

**Extended Data Figure 3.**
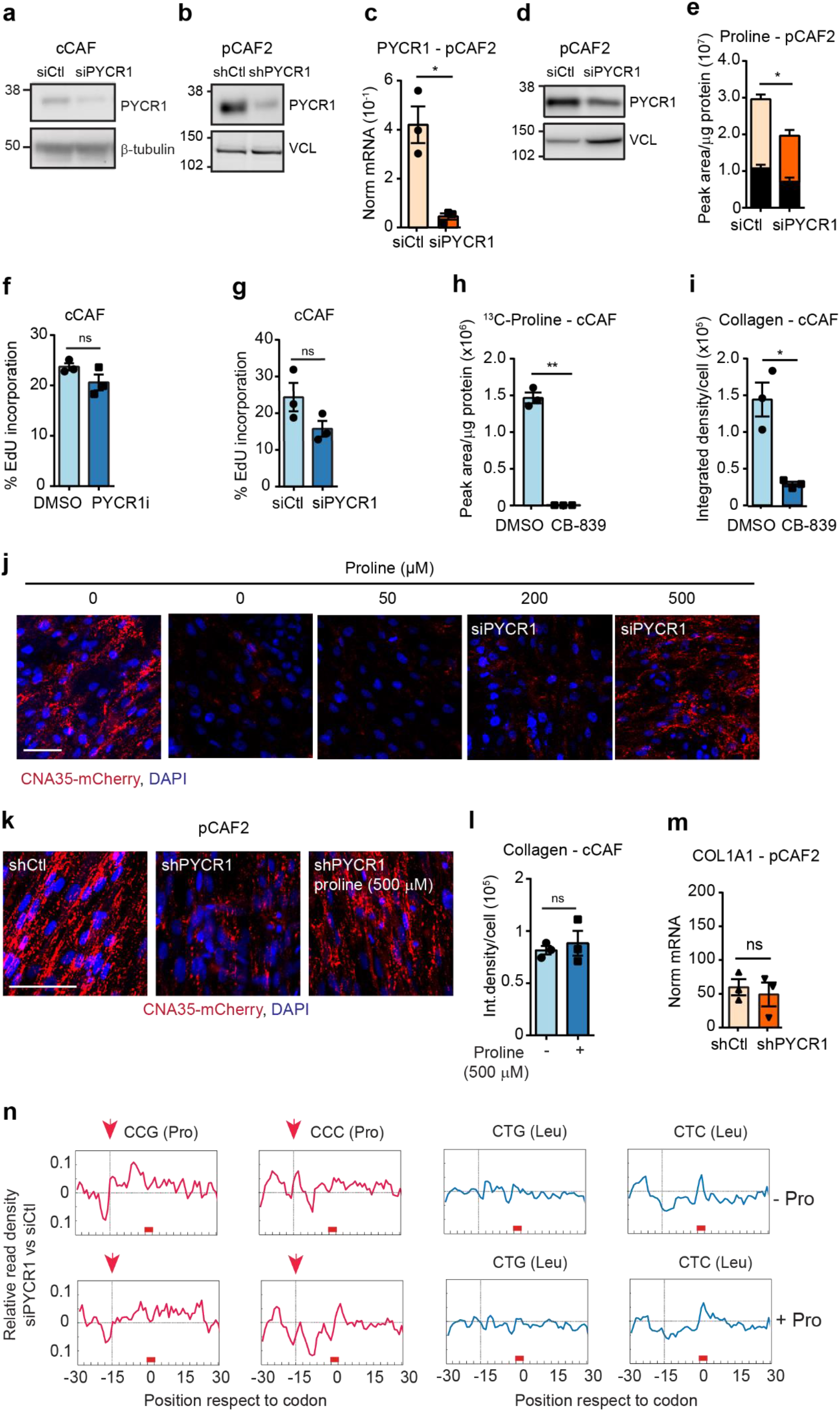
PYCR1 expression regulates proline synthesis and collagen production. **a.** Representative western blot of PYCR1 in cCAF transfected with siCtl or siPYCR1. β-tubulin was used as a loading control. **b.** Representative western blot of PYCR1 in shCtl and shPYCR1 pCAF2 lines. VCL = vinculin was used as a loading control. **c.** *PYCR1* expression in pCAF2 transfected with siCtl or siPYCR1, measured by RT-qPCR and normalised to 18S levels. N = 3 biological replicates. **d.** Representative western blot of PYCR1 in pCAF2 transfected with siCtl or siPYCR1. VCL = vinculin was used as a loading control. **e.** Total ^13^C-labelled (coloured bar) and unlabelled (black bar) proline in pCAF2 transfected with siCtl or siPYCR1 and labelled with ^13^C_5_-Glutamine, measured by MS-metabolomics. N = 3 biological replicates. **f.** Proliferation of cCAF treated with DMSO control or PYCR1i for 48h, measured by EdU incorporation. N = 3 biological replicates **g.** Proliferation of cCAF transfected with siCtl or siPYCR1, measured by EdU incorporation. N = 3 biological replicates. **h.** Total ^13^C-labelled proline measured by MS in cCAFs treated with 50 nM GLS inhibitor CB-839 or DMSO as control and labelled with ^13^C_5_-glutamine. N = 3 biological replicates. **i.** Quantification of collagen in cCAFs treated with 50 nM GLS inhibitor CB-839 or DMSO as control. N = 3 biological replicates. **j.** Representative image of CNA35-mCherry labelled collagen produced by cCAF siCtl or siPYCR1 treated with proline or PBS as control (quantification in Figure 3f). **k.** Representative image of CNA35-mCherry labelled collagen produced by pCAF shCtl or shPYCR1 treated with proline or PBS as control (quantification in Figure 3h). N = 4 biological replicates. **l.** Quantification of collagen produced by cCAF treated with proline or PBS as control. N = 3 biological replicates. Collagen was visualised with the collagen binding protein CNA35-mCherry. **m.** *COL1A1* expression in pCAF transfected with shCtl or shPYCR1, measured by RT-qPCR and normalised to 18S levels. N = 3 biological replicates. **n.** Ribosome read density, measured by Diricore analysis, at proline codons compared to Leucine codons in pCAF2 transfected with siCtl or siPYCR1. Error bars indicate mean ± SEM. *p ≤0.05, **p ≤0.01, ***p ≤0.001. Scale bar = 50 µm.

**Extended Data Figure 4.**
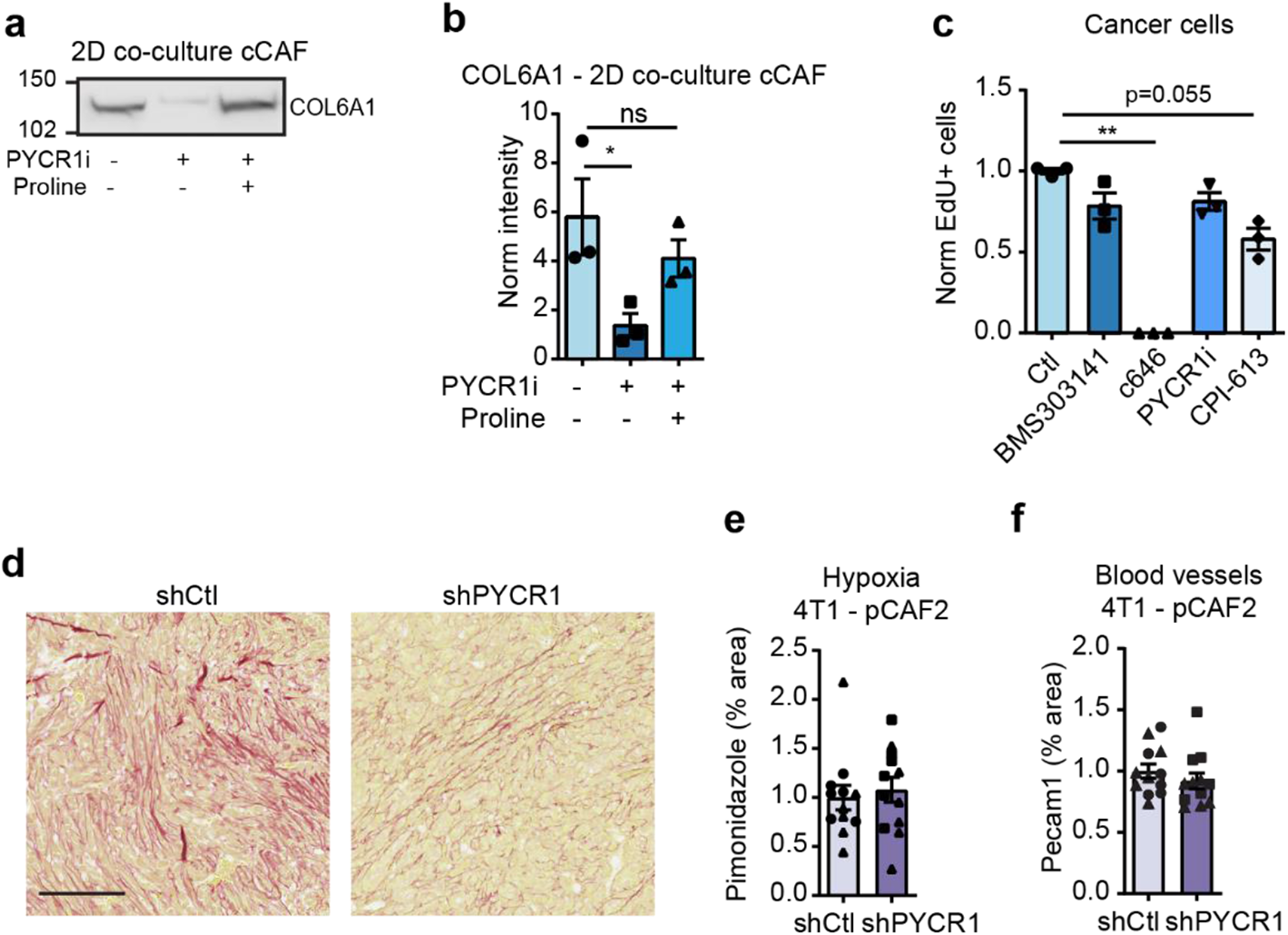
PYCR1 and COL1A1 expression in patient samples. **a, b.** Representative western blot (a) and quantification (b) of COL6A1 levels in decellularised ECM derived from co-cultures of cCAF and breast cancer cells treated with PYCR1i or DMSO, as control, and proline or PBS, as control. N = 3 biological replicates. **c.** Proliferation of breast cancer cells in mono-culture after 48h treatment with DMSO as control, c646, BMS303141, PYCR1i or CPI-613, measured by EdU incorporation. N = 3 biological replicates. **d.** Representative images of Siruis Red staining in FFPE sections of tumours from 4T1 cells expressing Aka-luciferase (AkaLuc) co-transplanted with pCAF2 shCtl or shPYCR1 (quantification in Figure 4s). **e.** Quantification of Pimonidazole staining in FFPE sections of tumours from shCtl/shPYCR1 CAF xenografts. N = 12 mice for each condition from two independent experiments. **f.** Quantification of Pecam1 staining in FFPE sections of tumours from 4T1 cells expressing Aka-luciferase (AkaLuc) co-transplanted with pCAF2 shCtl or shPYCR1. N = 12 mice for each condition from two independent experiments. Error bars indicate mean ± SEM. *p ≤0.05, **p ≤0.01, ***p ≤0.001. See Extended Data Figure 9 for red Ponceau staining of the blots used for COL6A1 staining in the ECM, which was used to normalise to total protein content in each lane.

**Extended Data Figure 5.**
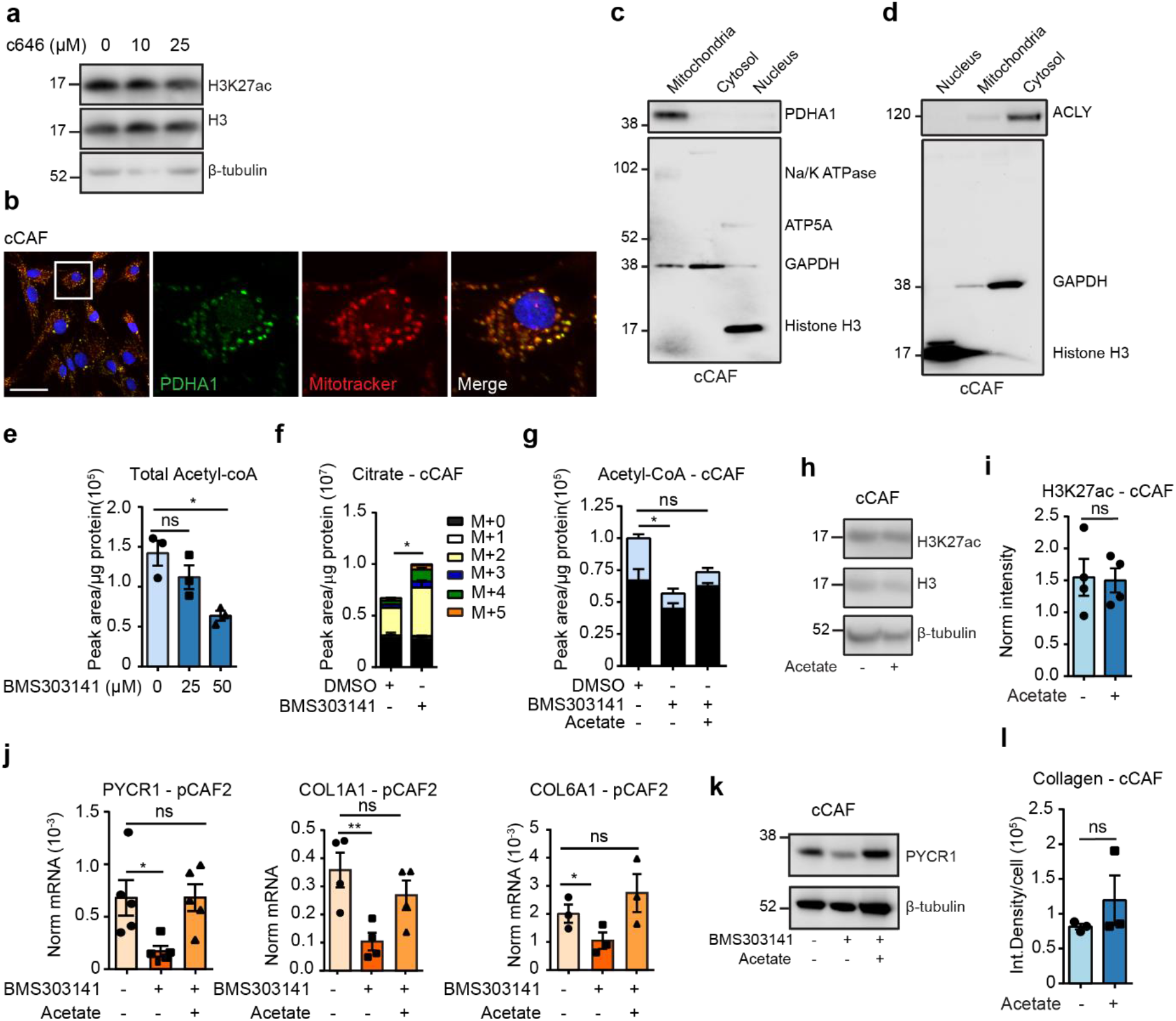
ACLY regulates acetyl-CoA production in CAFs. **a.** Representative western blot for histone H3 and H3K27ac of cCAFs treated with increasing doses of the EP300 inhibitor c646. β-tubulin was used as loading control. **b.** Representative confocal microscopy images of cCAFs showing co-localisation of PDHA1 with mitochondria (MitoTracker). Scale bar = 50 µm. **c.** Western blot of PDHA1 in mitochondrial, nuclear and cytosolic fractions of cCAFs. Na/K ATPase = membrane marker, ATP5A = mitochondrial marker, GAPDH = cytosolic marker, H3 = nuclear marker. **d.** Western blot of ACLY in mitochondrial, nuclear and cytosolic fractions of cCAFs. ATPase = membrane marker, ATP5A = mitochondrial marker, GAPDH = cytosolic marker, H3 = nuclear marker. **e**. Total acetyl-CoA in cCAF treated with DMSO, as control, or BMS303141 at the indicated concentrations, measured by MS-metabolomics. N = 3 biological replicates. **f.** Total levels of ^13^C-glucose labelled (coloured) and unlabelled (black) citrate in cCAFs treated with DMSO as control or BMS303141, measured by MS-metabolomics. N = 3 biological replicates. **g.** Total ^13^C-labelled (coloured bar) and unlabelled (black bar) acetyl-CoA in cCAF treated with DMSO, as control, or BMS303141 and acetate or PBS, as control, and labelled with ^13^C_6_-Glucose, measured by MS-metabolomics. N = 3 biological replicates. **h,i.** Representative western blot (h) and quantification (i) of H3K27 levels in cCAFs ± 1 mM acetate or PBS as control. β-tubulin was used as loading control. N = 3 biological replicates. **j.** mRNA expression of *PYCR1, COL1A1 and COL6A1* in pCAF2 treated with DMSO, as control, or BMS303141 and acetate or PBS, as control, measured by RT-qPCR and normalised to 18S expression. N = 3-5 biological replicates. **k.** Representative western blot for PYCR1 in cCAF treated with DMSO, as control, or BMS303141 and acetate or PBS, as control. **l.** Quantification of collagen produced by cCAFs treated with acetate or PBS as control. N = 3 biological replicates. Error bars indicate mean ± SEM. *p ≤0.05, **p ≤0.01, ***p ≤0.001.

**Extended Data Figure 6.**
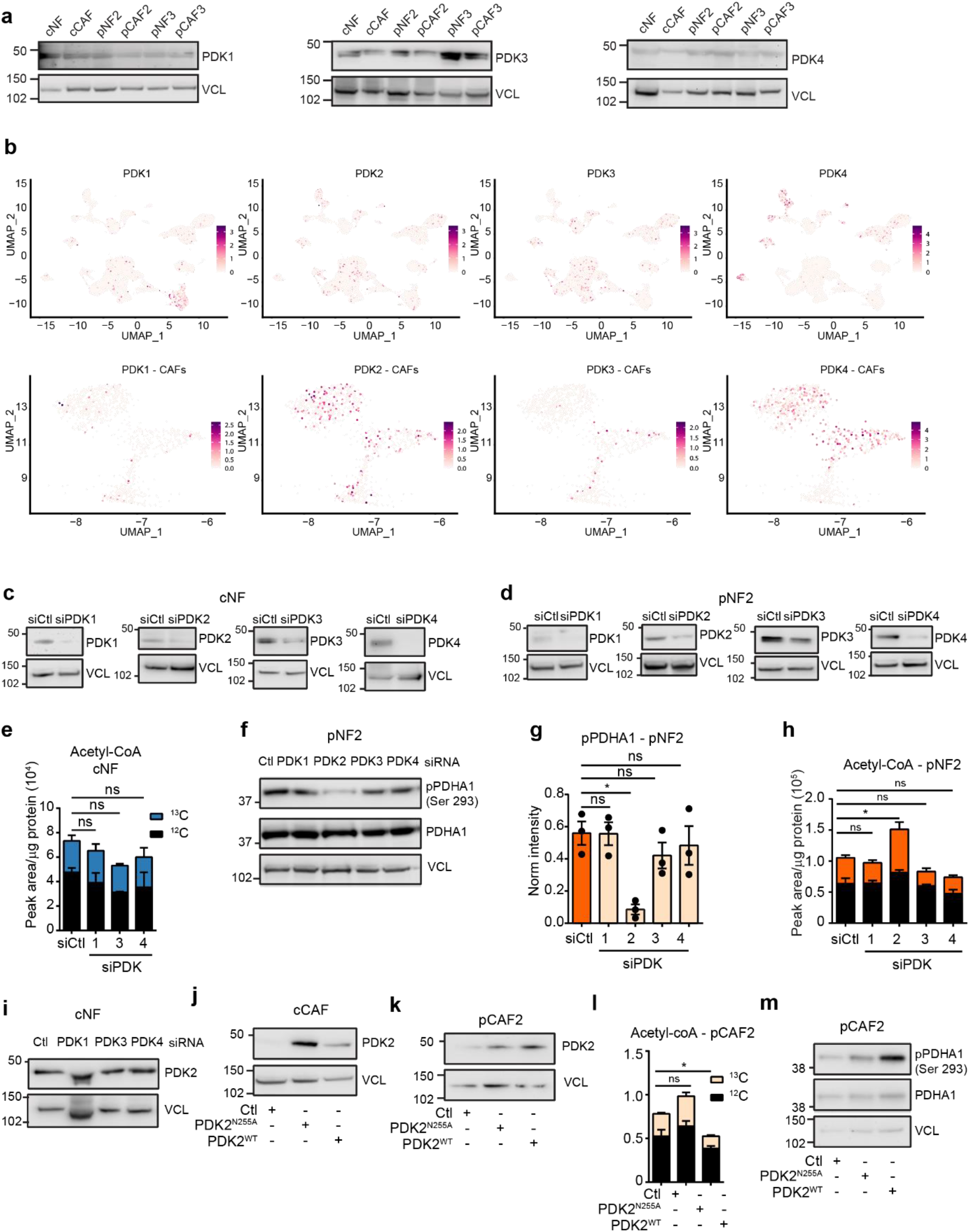
PDK2 regulates acetyl-coA production in patient derived fibroblasts. **a.** Representative western blots showing levels of PDK1, PDK3 and PDK4 in mammary NFs and CAFs. VCL = vinculin was used as a loading control. **b**. UMAP visualisation of stromal, immune and cancer cells (top) or CAF-only (bottom) aligned using canonical correlation analysis in Seurat. Cell types defined as in Wu et al. **c,d.** Representative western blots showing expression of PDK1-4 after transfection with siCtl or the respective siRNA in cNF (c) and pNF2 (d). VCL = vinculin was used as a loading control. **e.** ^13^C-labelled (coloured bar) and unlabelled (black bar) acetyl-CoA measured by MS in cNF2 transfected with siCtl or siPDK1,3,4 and labelled with ^13^C_6_-glucose. N = 3 biological replicates. **f,g.** Representative western blot (f) and quantification (g) showing pPDHA1 levels in pNF2 transfected with siCtl or siPDK1-4. N = 3 biological replicates. Total PDHA1 and VCL (vinculin) were used as a loading control. **h.** ^13^C-labelled (coloured bar) and unlabelled (black bar) acetyl-CoA measured by MS in pNF2 transfected with siCtl or siPDK1-4 and labelled with ^13^C_6_-glucose. N = 3 biological replicates. **i.** Representative Western blot showing PDK2 levels in cNF transfected with siCtl, siPDK1, siPDK3, or siPDK4. VCL = Vinculin was used as a loading control. **j,k.** Representative Western blot showing the levels of overexpression of PDK2 wild type (PDK2^WT^) or the enzymatically inactive mutant form (PDK2^N255A^) in cCAF (j) and pCAF2 (k) after transfection with empty vector (Ctl), pGC-PDK2^N255A^ or pGC-PDK2^WT^. VCL = vinculin was used as a loading control. **l.** ^13^C-labelled (coloured bar) and unlabelled (black bar) acetyl-CoA measured by MS in pCAF2 transfected with empty vector, pGC-PDK2^N255A^ or pGC-PDK2^WT^ and labelled with ^13^C_6_-glucose. N = 3 biological replicates. **m.** Representative western blot showing PDHA1 phosphorylation levels at serine 293 in pCAF2 after transfection with empty vector, pGC-PDK2^N255A^ or pGC-PDK2^WT^. VCL = vinculin was used as a loading control. Error bars indicate mean ± SEM. *p ≤0.05.

**Extended Data Figure 7.**
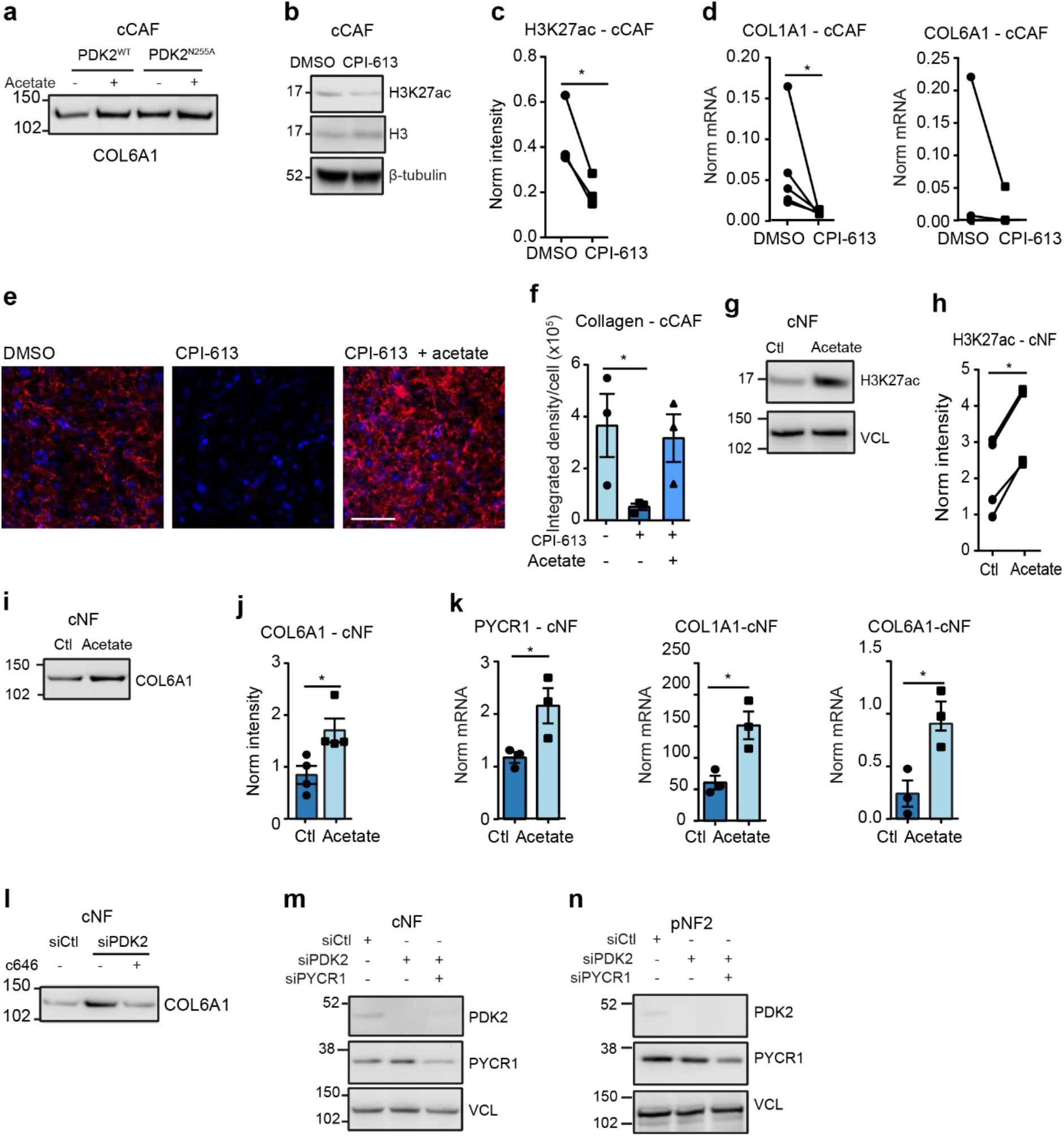
PDH inhibition regulates collagen production. **a.** Representative western blot showing COL6A1 levels in ECM derived from cCAF transfected with pGC-PDK2^N255A^ or pGC-PDK2^WT^ and treated with acetate or PBS as control. **b, c.** Representative western blot (b) and quantification (c) of H3K27ac levels in cCAF treated with DMSO as control or CPI-613. β-tubulin was used as a loading control. N = 3 biological replicates. **d.** Expression of *COL1A1 and COL6A1* in cCAF treated with DMSO as control or CPI-613, measured by RT-qPCR and normalised to 18S expression. N = 3 biological replicates. **e, f.** Representative confocal microscopy images (e) and quantification (f) of collagen produced by cCAF treated with DMSO, as control, or CPI-613 and acetate or PBS, as control. N = 3 biological replicates. Scale bar = 100 µm. **g, h.** Representative western blot (g) and quantification (h) of H3K27ac levels in cNF treated with acetate or PBS as control. VCL = vinculin was used as a loading control. N = 3 biological replicates. **i, j.** Representative western blot (i) and quantification (j) of COL6A1 levels in decellularised ECM derived from cNF treated with acetate or PBS as control. N = 3 biological replicates. **k.** Expression of *PYCR1, COL1A1 and COL6A1* in cNF treated with acetate or PBS as control, measured by RT-qPCR and normalised to 18S expression. N = 3 biological replicates. **l.** Representative western blot of COL6A1 levels in decellularised ECM derived from cNF transfected with siCtl or siPDK2 and treated with DMSO as control or c646. **m, n**. Representative western blots showing PDK2 and PYCR1 levels in cNF (m) and pNF2 (n) transfected with siCtl, siPDK2 or siPDK2 and siPYCR1. VCL = vinculin was used as a loading control. Error bars indicate mean ± SEM. *p ≤0.05, **p ≤0.01, ***p ≤0.001. Scale bar = 50 µm. See Extended Data Figure 9 for red Ponceau staining of the blots used for COL6A1 staining in the ECM, which was used to normalise to total protein content in each lane.

**Extended Data Figure 8.**
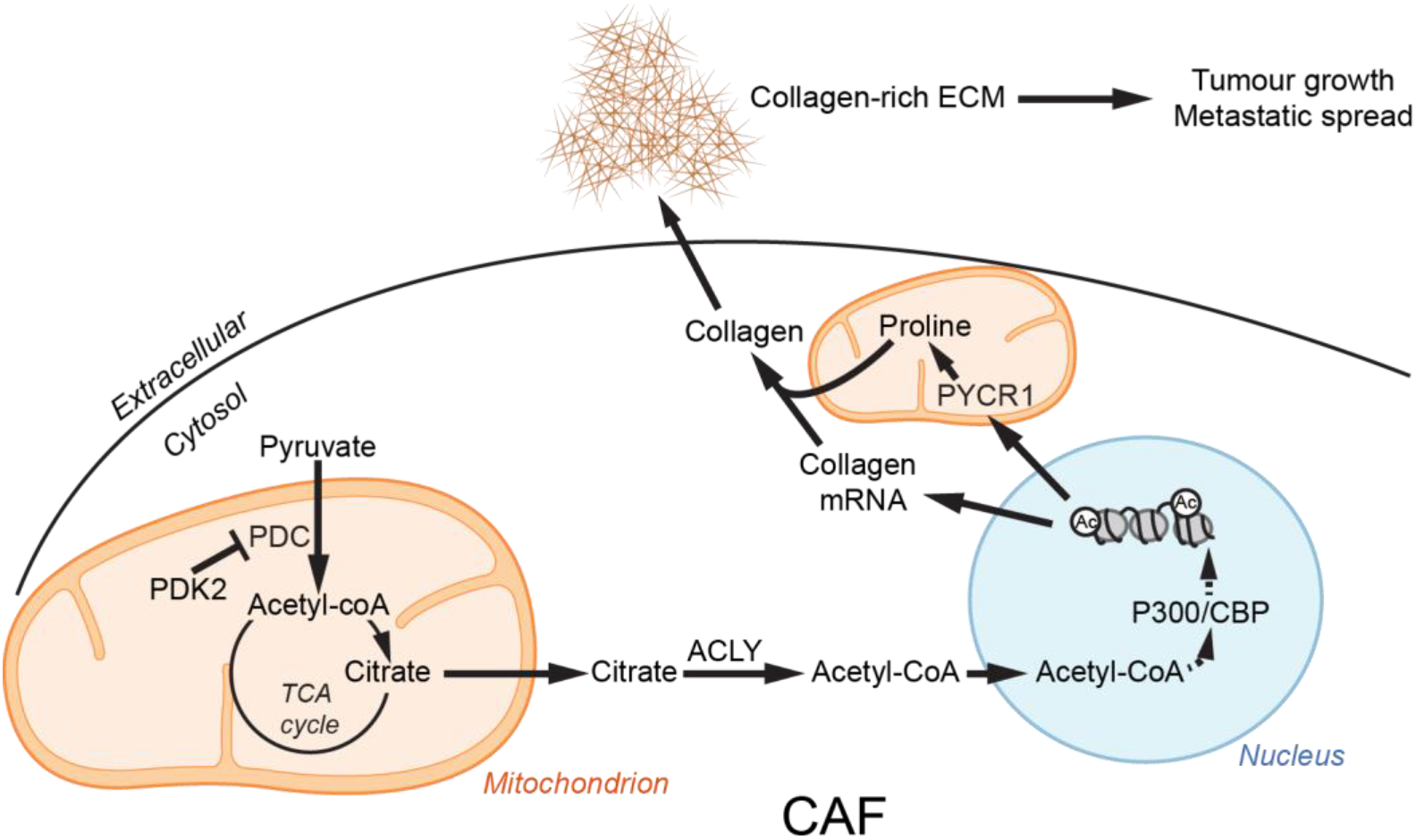
Working model. Scheme showing how PDH-derived acetyl-CoA and PYCR1-derived proline act together to stimulate and maintain collagen production in CAFs.

**Extended Data Figure 9.**
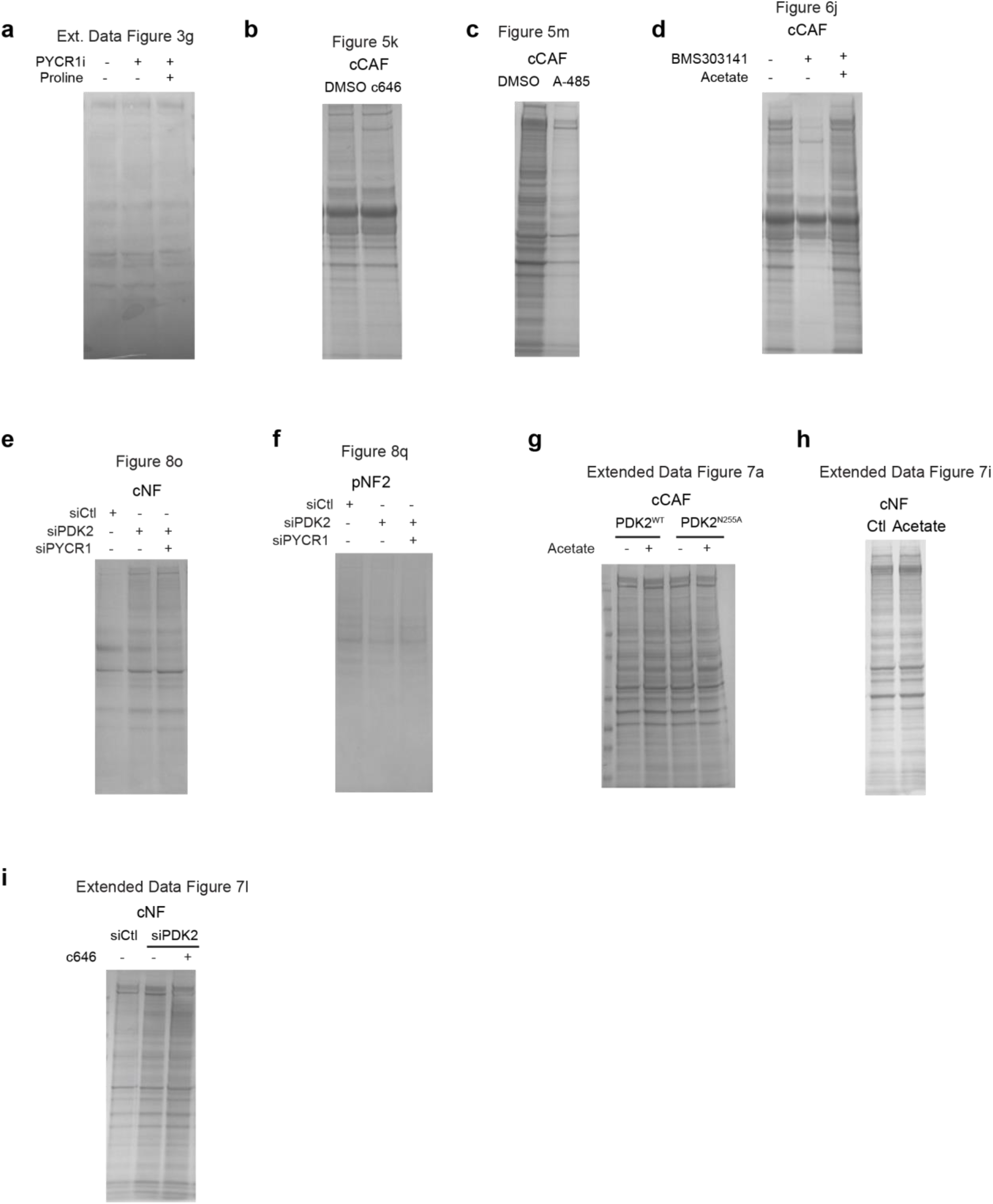
Loading controls for ECM western blots. **a.** Ponceau S total protein stain for blot shown in Figure 3g. **b.** Ponceau S total protein stain for blot shown in Figure 5k. **c.** Ponceau S total protein stain for blot shown in Figure 5m. **d.** Ponceau S total protein stain for blot shown in Figure 6j. **e.** Ponceau S total protein stain for blot shown in Figure 8o. **f.** Ponceau S total protein stain for blot shown in Figure 8q. **g.** Ponceau S total protein stain for blot shown in Extended Data Figure 7a. **h.** Ponceau S total protein stain for blot shown in Extended Data Figure 7i. **i.** Ponceau S total protein stain for blot shown in Extended Data Figure 7l.

## Acknowledgements

We thank Leo Carlin, Frederic Fercoq, Ewan McGhee, Lynn Mcgarry and the Cancer Research UK Beatson Institute core research services and advanced technology facilities, including BSU, HiCAR, BAIR, and histology; Clare Orange for histopathology services; NHS Greater Glasgow and Clyde Biorepository for providing patient samples, and the PRIDE team. The results shown in Extended Figure 4h are based upon data generated by the TCGA Research Network: https://www.cancer.gov/tcga.

This work was funded by Cancer Research UK (CRUK Beatson Institute A31287, CRUK Glasgow Centre A18076, and Stand Up to Cancer campaign for Cancer Research UK A29800 (to S.Z.)) and Breast Cancer Now (2019AugPR1307 to S.Z.). E.G. was supported by the European Union’s H2020 programme (675585 Marie-Curie ITN ‘‘SymBioSys’’) and JRC for Computational Biomedicine, which is partially funded by Bayer, K.P. was supported by AMS Biotechnology (Europe) Ltd and the University of Strathclyde. M.M. was supported by an ERC Consolidator grant (ImmunoFit).

## Author contributions

Conceptualization: E.J.K. and S.Z.; methodology: E.J.K., D.S., M.Z., S.L., K.B., F.L-P., J.J.K. J.S-R., C.M., H.D., S.T., M.M., K.K. and S.Z.; investigation: E.J.K., K.P., E.S., C.B., L.J.N., J.R.H-F., S.L., G.K., S.D., A.H., E.G., C.R.D. and S.Z.; resources: C.J., R.M.J., R.S., G.M. and M.P.; writing original draft, review and editing: E.J.K., K.P, M.Z., M.M., S.T. H.D. and S.Z.; supervision: M.P., C.M., J.J.K., F.L-P., J.S-R., K.B., M.M., M.Z. and S.Z.

## Declaration of Interests

We declare no competing interests.

## Notes

### Competing Interest Statement

The authors have declared no competing interest.

